# The Influence of P_₂_O_₅_ on the Structure, Crystallization, and Bioactivity of Silicate-Based Bioactive Glasses and Glass-Ceramics

**DOI:** 10.1101/2025.03.30.644796

**Authors:** Mahmoud G. Soliman, M. I. EL Gohary

**Author notes:** Corresponding author: M. G. Soliman.

## Abstract

This study investigates the effects of P_2_O_5_ addition on the structural, thermal, and bioactive properties of silicate-based bioactive glasses and glass-ceramics. Four glass compositions (G0, G1, G2, G3) with varying P_2_O_5_ content (0.0, 2.5, 5.0, and 7.5 wt%) were prepared using the melt-quenching method. The samples were characterized using differential thermal analysis, X-ray diffraction, scanning electron microscopy, Fourier-transform infrared spectroscopy, and in vitro bioactivity tests in simulated body fluid. Results indicate that P_2_O_5_ addition lowers the glass transition temperature, promotes phase separation, and enhances the crystallization of fluoroapatite while reducing the formation of fluorophlogopite. *In vitro* tests revealed that P_2_O_5_ accelerates the formation of a hydroxycarbonate apatite layer on the glass surface, indicating improved bioactivity. However, in glass-ceramics, P_2_O_5_ delays ion release and HCA formation due to the stabilization of the residual glass phase. These findings suggest that P_2_O_5_ plays a critical role in tailoring the bioactivity and mechanical properties of bioactive glasses and glass-ceramics for biomedical applications.

## 1. Introduction

Bioactive glasses and glass-ceramics have gained significant attention in biomaterials research due to their unique ability to bond with living tissues through the formation of a hydroxycarbonate apatite (HCA) layer.^1-4^ This property makes them highly suitable for applications in bone repair, dental implants, and tissue engineering. Hench first introduced the concept of bioactive materials in the 1970s with the development of the 45S5 Bioglass®, which demonstrated the ability to form a strong bond with bone tissue.^5^ Since then, extensive research has been conducted to optimize the composition, structure, and properties of bioactive glasses to enhance their performance in biomedical applications.

The bioactivity of these materials is primarily attributed to their surface reactivity in physiological environments. When exposed to body fluids, bioactive glasses undergo a series of surface reactions, including ion exchange, network dissolution, and precipitation, leading to the formation of an HCA layer.^6^ This layer is chemically and structurally similar to the mineral phase of bone, facilitating the integration of the implant with the surrounding tissue. The rate and extent of HCA formation depend on the glass composition, with key components such as SiO_2_, CaO, Na_2_O, and P_2_O_5_ playing critical roles in determining the bioactivity.^6, 7^

Among these components, phosphorus pentoxide (P_2_O_5_) has been identified as a crucial element in enhancing the bioactivity of silicate-based glasses. P_2_O_5_ promotes the formation of apatite phases, which are essential for bone bonding^8, 9^. However, its role in the crystallization behavior and overall bioactivity of glass-ceramics is complex and not fully understood. For instance, while P_2_O_5_ accelerates apatite crystallization in glasses, it can also stabilize the residual glass phase in glass-ceramics, delaying ion release and HCA formation.^10, 11^ Additionally, P_2_O_5_ influences the mechanical properties of these materials by altering phase composition, such as suppressing fluorophlogopite crystallization in favor of fluoroapatite.^12^

Despite extensive research, the interplay between P_2_O_5_ content, crystallization kinetics, and bioactivity remains incompletely understood. For example, Kokubo et al.^9^ demonstrated that CaO·SiO_2_ glasses free of P_2_O_5_ could still form HCA layers, while Marghussian et al.^13^ suggested that reducing P_2_O_5_ content might enhance mechanical properties without compromising bioactivity. These findings highlight the need for systematic studies to elucidate the optimal P_2_O_5_ concentrations for specific applications.

Understanding the effects of P_2_O_5_ on the structure, crystallization, and bioactivity of silicate-based bioactive glasses and glass-ceramics is essential for designing materials with optimized properties for specific biomedical applications. This study aims to systematically investigate the influence of P_2_O_5_ addition on the thermal, structural, and bioactive properties of silicate-based glasses and glass-ceramics. Four glass compositions with varying P_2_O_5_ content (0.0, 2.5, 5.0, and 7.5 wt%) were prepared, and their properties were characterized using differential thermal analysis (DTA), X-ray diffraction (XRD), scanning electron microscopy (SEM), Fourier-transform infrared spectroscopy (FTIR), and *in vitro* bioactivity tests in simulated body fluid (SBF).

The results of this study provide valuable insights into the role of P_2_O_5_ in tailoring the properties of bioactive glasses and glass-ceramics. By understanding the effects of P_2_O_5_ on the crystallization behavior and bioactivity of these materials, it is possible to design compositions with enhanced performance for bone repair and tissue engineering applications. This research contributes to the ongoing efforts to develop advanced biomaterials that can improve the outcomes of medical treatments and enhance the quality of life for patients.

## 2. Materials and Methods

### 2.1 Glass Preparation

Four glass compositions based on the SiO_2_-CaO-MgO-MgF_2_-K_2_O system with varying P_2_O_5_ content (0.0, 2.5, 5.0, and 7.5 wt%) were prepared using the melt-quenching method.^14-17^ The raw materials, including quartz (SiO_2_), calcium carbonate (CaCO_3_), magnesium carbonate (MgCO_3_), magnesium fluoride (MgF_2_), potassium carbonate (K_2_CO_3_), and phosphorus pentoxide (P_2_O_5_), were weighed in appropriate proportions to achieve the desired compositions. The batches were thoroughly mixed in an agate ball mill for 30 minutes to ensure homogeneity. The mixed batches were then preheated in a platinum crucible at 1050°C for 1 h to remove volatile components and decarbonize the carbonates. After preheating, the batches were melted in a platinum crucible at 1450°C for 2 h in an electric furnace. The molten glass was quenched in deionized water to obtain glass frits. The glass frits were dried at 100°C for 12 h and then ground into fine powders using an agate mortar and pestle. The powders were sieved to obtain particle sizes of 50-80 µm,^17^ which were used for subsequent characterization and *in vitro* bioactivity tests.

### 2.2 Thermal Analysis

Differential thermal analysis (DTA) was performed to determine the glass transition temperature (Tg), crystallization temperature (Tc), and melting temperature (Tm) of the glasses.^18^ The glass powders were heated in a DTA instrument (Labsys™ TG-DSC16) at a rate of 10°C/min in an argon atmosphere, with α-Al_2_O_3_ used as a reference material. The DTA curves were recorded, and the characteristic temperatures were determined from the onset and peak positions of the endothermic and exothermic effects^6^.

### 2.3 Heat Treatment and Crystallization

Based on the DTA results, the glass samples were subjected to a two-step heat treatment to induce crystallization and obtain glass-ceramics. The first step involved nucleation at a temperature close to the glass transition temperature (Tg), and the second step involved crystal growth at a higher temperature (Tc). The glasses were heat-treated at 750°C and 900°C for 2 h in a muffle furnace. The heat-treated samples were cooled to room temperature in the furnace to avoid thermal shock and cracking^8^.

### 2.4 Structural Characterization

Equation 1 evaluated the degree of crystallinity, corresponding to the fraction of crystalline phase present in the examined sample.^19^

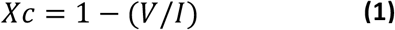

Where I is the intensity of the main peak reflection, and V is the intensity of the hollow between the main peak and the peak beside it.

The crystalline phases in the glasses and glass-ceramics were identified using XRD, which were recorded using a Diano Corporation diffractometer with CoKα radiation (λ = 1.54 Å) in the 2θ range of 10° to 80° with a step size of 0.1° and a counting time of 1 second per step. The degree of crystallinity and crystal size were calculated using Scherrer’s equation– Equation 2.^19^

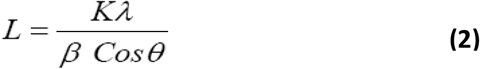

Where L is the crystallite size, K is the Scherrer constant (0.9), λ is the wavelength of the X-ray radiation, β is the full width at half maximum (FWHM) of the diffraction peak, and θ is the diffraction angle.

The microstructure of the glass-ceramics was examined using scanning electron microscopy (SEM). The samples were coated with a thin layer of gold using a sputter coater (Edwards 5150) to enhance conductivity.^10^ SEM images were acquired using a Philips XL30 microscope at an accelerating voltage of 30 kV and magnifications ranging from 500x to 2000x.

### 2.5 Density and Porosity Measurements

The density of the glasses and glass-ceramics was measured using the Archimedean method.^18^ The samples were weighed in air (W) and in deionized water (W_w_), and the density was calculated using Equation 3.

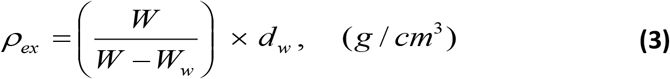

where ρ_ex_ is the experimental density, W is the weight of the sample in air, W_w_ is the weight of the sample in water, and *d*_w_ is the density of water (1 g/cm^3^). For glass-ceramics, the samples were coated with paraffin wax to prevent water absorption, and the density was calculated using a modified equation 4 to account for the wax coating^11^.

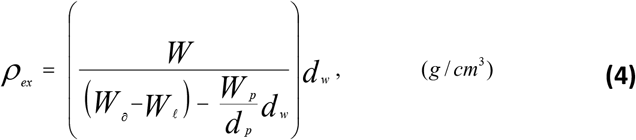

Experimental densities were obtained by Archimedes’ methods in two ways: The first, the pellet was sealed with paraffin wax to prevent water absorption. In the second way, the pellet was immersed in de-ionized water for 20 h to fill the inner pores with water.

Where ρ_ex_ is the experimental (bulk) density of the sample, W is the weight of the sample in air, W_a_ is the weight of the sample coated with paraffin wax in air, W_*l*_ is the weight of sample coated with paraffin wax in water, W_o_ is the weight of the thin layer of paraffin wax coat, W_w_ = (W_a_ - W), *d*_*p*_ is the density of paraffin wax, (*d*_*p*_ = 0.79 g/cm^3^ at RT), and *d*_*w*_ is the density of de-ionized water, (*d*_w_ = 1 gm/cm^3^ at RT).

The porosity of the glass-ceramics was determined by measuring the water absorption.^20^ The samples were soaked in deionized water for 48 h, and the wet weight (W_wet), dry weight (W_dry), and suspended weight (W_suspended) were measured. The apparent porosity was calculated using Equations 5 & 6.

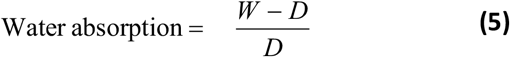

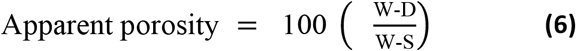

Where W is the wet weight, D is the dry weight, and S is the wet weight suspended in water

### 2.6 Microhardness Measurements

The microhardness of the glass-ceramics was measured using a Vickers microhardness tester (Shimadzu HMV-2).^12, 21^ Indentations were made on the surface of the samples using a square-based pyramidal diamond indenter with a face angle of 136°. A load of 25 kg was applied for 15 sec., and the diagonal lengths of the indentations were measured using a calibrated microscope. The Vickers microhardness (Hv) was calculated using Equation 7.

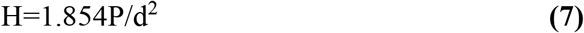

Where P is the applied load (kg), d is the diagonal length of the indentation (mm). Five indentations were made on each specimen with a constant load of 25 Kg/15 sec.

### 2.7 *In Vitro* Bioactivity Tests

The bioactivity of the glasses and glass-ceramics was evaluated by immersing the samples in simulated body fluid (SBF) at 37°C for 30 days.^6, 22^ The SBF was prepared according to the protocol described by Kokubo et al.,^6^ with ion concentrations nearly equal to those of human blood plasma. The samples were immersed in SBF at a powder-to-solution ratio of 0.25 g/100 mL. The pH of the SBF was monitored using a pH meter (Jenway 3510), and the concentrations of Ca^2+^ and P^5+^ ions in the SBF were measured using a UV-Vis spectrophotometer (Jenway 4600) with biological kits.

After immersion, the samples were removed from the SBF, rinsed with acetone, and dried at room temperature. The formation of an HCA layer on the surface of the samples was analyzed using FTIR spectroscopy (Perkin-Elmer Model 580) in the wave number range of 4000-400 cm^-1^. The FTIR spectra were recorded with a resolution of 4 cm^-1^, and the functional groups were identified based on previous assignments in the literature.^7^

## 3. Results and Discussion

### 3.1 Glass Formation and Morphology

DTA revealed significant changes in the thermal properties of the glasses with increasing P_2_O_5_ content, as shown in Figure 1 and Table S1. The glass transition temperature (Tg) decreased from 618°C for G0 to 595°C for G3, indicating that P_2_O_5_ weakens the silicate network and reduces the viscosity of the glass. This reduction in Tg is attributed to the disruption of the Si-O-Si network by P_2_O_5_, which introduces non-bridging oxygen (NBO) atoms and lowers the overall connectivity of the glass structure.^12^ The crystallization temperature (Tc) also decreased with increasing P_2_O_5_ content, from 881°C for G0 to 750°C for G3, suggesting that P_2_O_5_ promotes early crystallization.^23^ The exothermic peaks in the DTA curves were attributed to the crystallization of fluoroapatite, wollastonite (CaSiO_3_), and fluorophlogopite (KMg_2_._5_Si_4_O_10_F_2_). The broadening of the exothermic peaks in G1, G2, and G3 indicates that P_2_O_5_ addition leads to a wider range of crystallization temperatures, likely due to the formation of multiple crystalline phases.^10^

**Figure 1.**
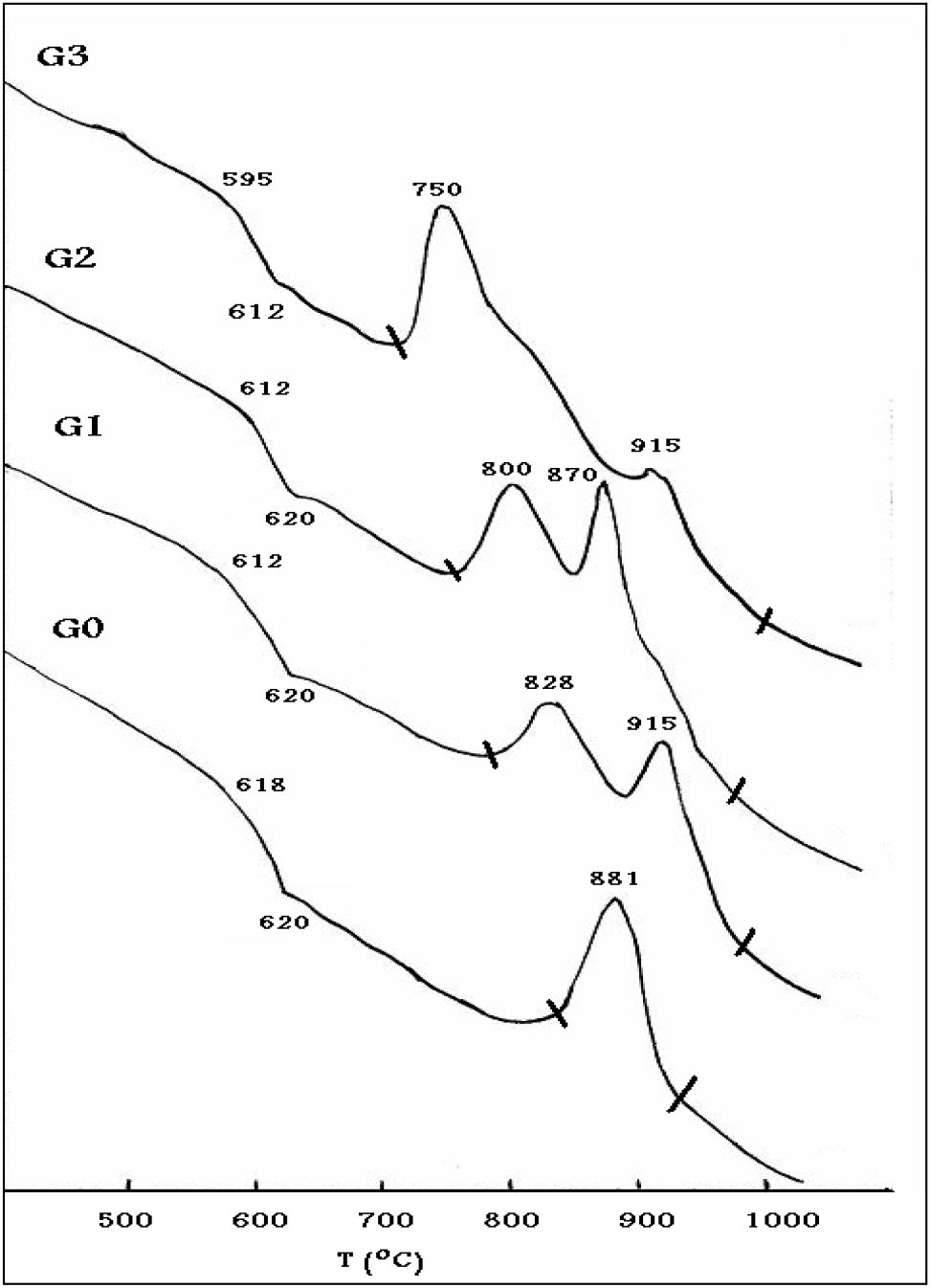
DTA curves of the investigated glasses

The base glass (G0) and the glass with 2.5 wt% P_2_O_5_ (G1) were transparent, indicating a homogeneous amorphous structure. In contrast, the glasses with higher P_2_O_5_ content (G2 and G3) exhibited opalescence and faint white coloration, suggesting the onset of phase separation and partial crystallization. This visual observation was corroborated by XRD analysis as can be seen in Figures 2-4 and Table S2, which confirmed the amorphous nature of G0 and G1, while G2 and G3 showed minor crystalline phases of forsterite (Mg_2_SiO4), cristobalite (SiO_2_), and fluoroapatite (Ca_5_(PO4)_3_F).^23^ The presence of these crystalline phases in G2 and G3 indicates that P_2_O_5_ promotes phase separation and nucleation, even during the cooling process of the melt (Figure 2). This phenomenon is consistent with previous studies, which have shown that P_2_O_5_ can act as a nucleating agent, facilitating the crystallization of specific phases in silicate glasses.^8, 24^ XRD reference patterns are provided in the Supporting Information (Figures S1-S8).

**Figure 2.**
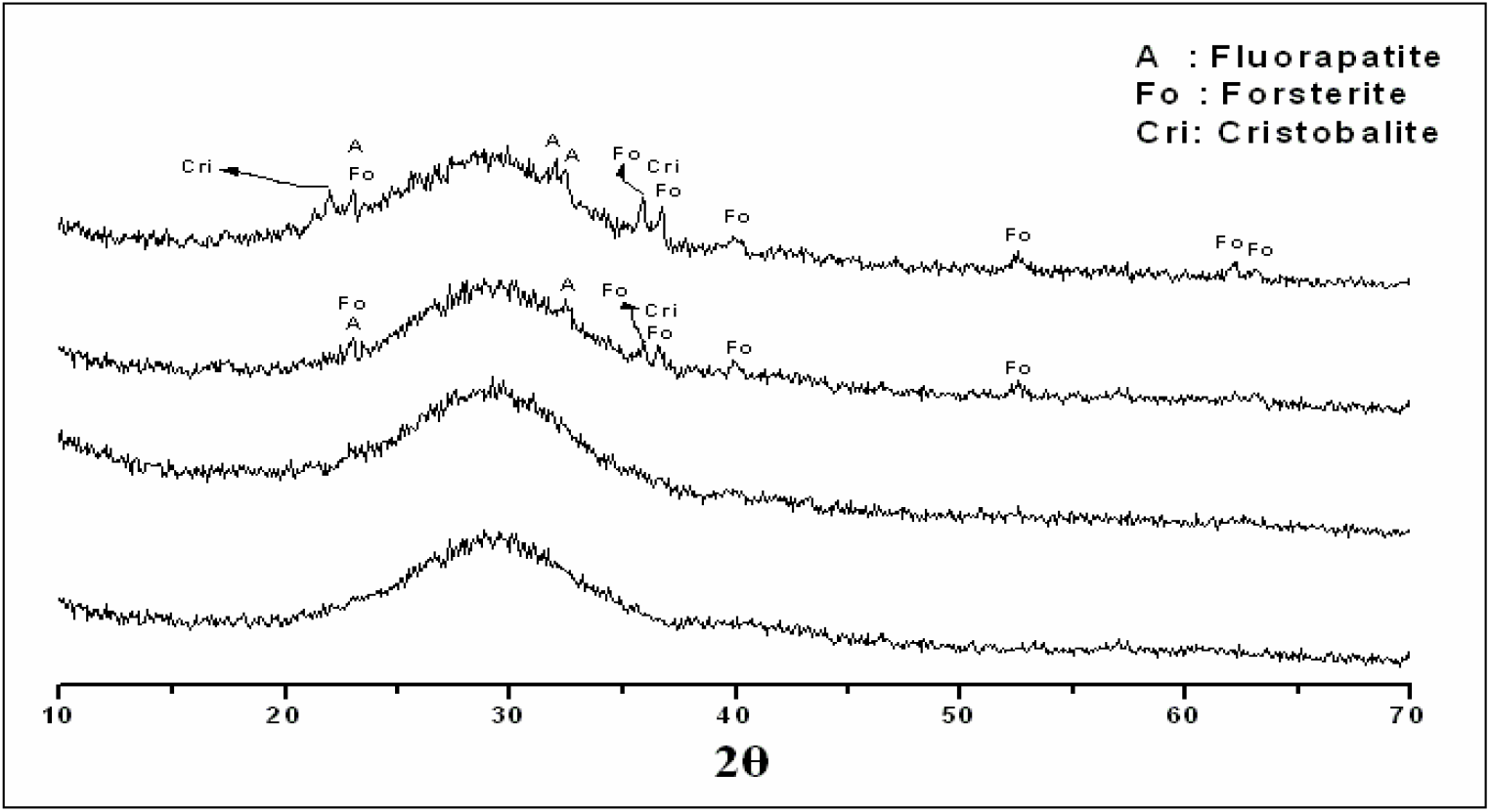
XRD patterns of the glass samples G0, G1, G2, and G3.

Figure 3 presents the XRD patterns and the crystalline phases precipitated in the G0, G1, G2, and G3 samples after crystallization at 750 °C for 2 hours. The G0 parent glass remained fully amorphous, with no detectable crystalline phases. However, upon doping with 2.5 wt% P_2_O_5_, minor precipitation of fluorophlogopite and fluoroapatite was observed. A further increase in P_2_O_5_ content led to the formation of highly crystalline materials, as evidenced by the more pronounced diffraction peaks in G2 and G3 compared to G1 (Figure 3).

**Figure 3.**
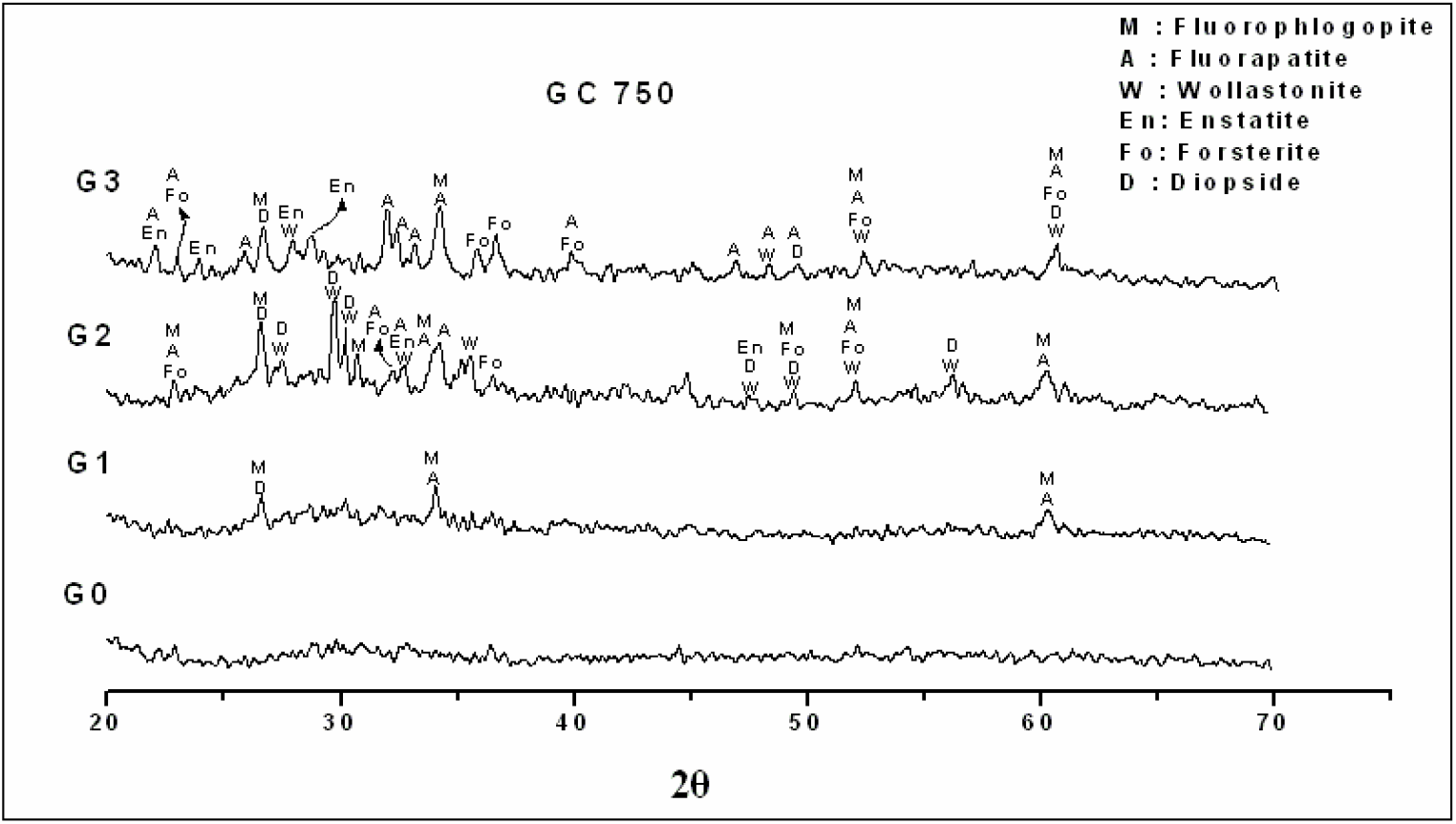
XRD patterns of the glass samples G0, G1, G2, and G3 crystallized at 750 °C for 2 h.

However, an increase in heat treatment up to 900°C resulted in the formation of highly crystalline glass-ceramics. XRD analysis (Figure 4) showed that the addition of P_2_O_5_ significantly influenced the crystallization behavior of the glasses. In G0, the primary crystalline phase was fluorophlogopite, with minor amounts of wollastonite and diopside (CaMgSi_2_O6). With increasing P_2_O_5_ content, the crystallization of fluoroapatite became more pronounced, while the formation of fluorophlogopite was suppressed. This shift in crystallization behavior is attributed to the preferential association of Ca^2+^ and Mg^2+^ ions with phosphate-rich phases, which inhibits the crystallization of silicate-rich phases such as fluorophlogopite.^19^ SEM images revealed a dense microstructure with interlocked flakes of fluorophlogopite in G0, which gradually decreased with increasing P_2_O_5_ content. In contrast, the crystal size of fluoroapatite increased with P_2_O_5_ addition, indicating that P_2_O_5_ promotes the growth of apatite crystals.^12, 25^ The microstructural changes observed in the glass-ceramics are consistent with the XRD results and highlight the role of P_2_O_5_ in tailoring the crystallization behavior of silicate-based glasses.

**Figure 4.**
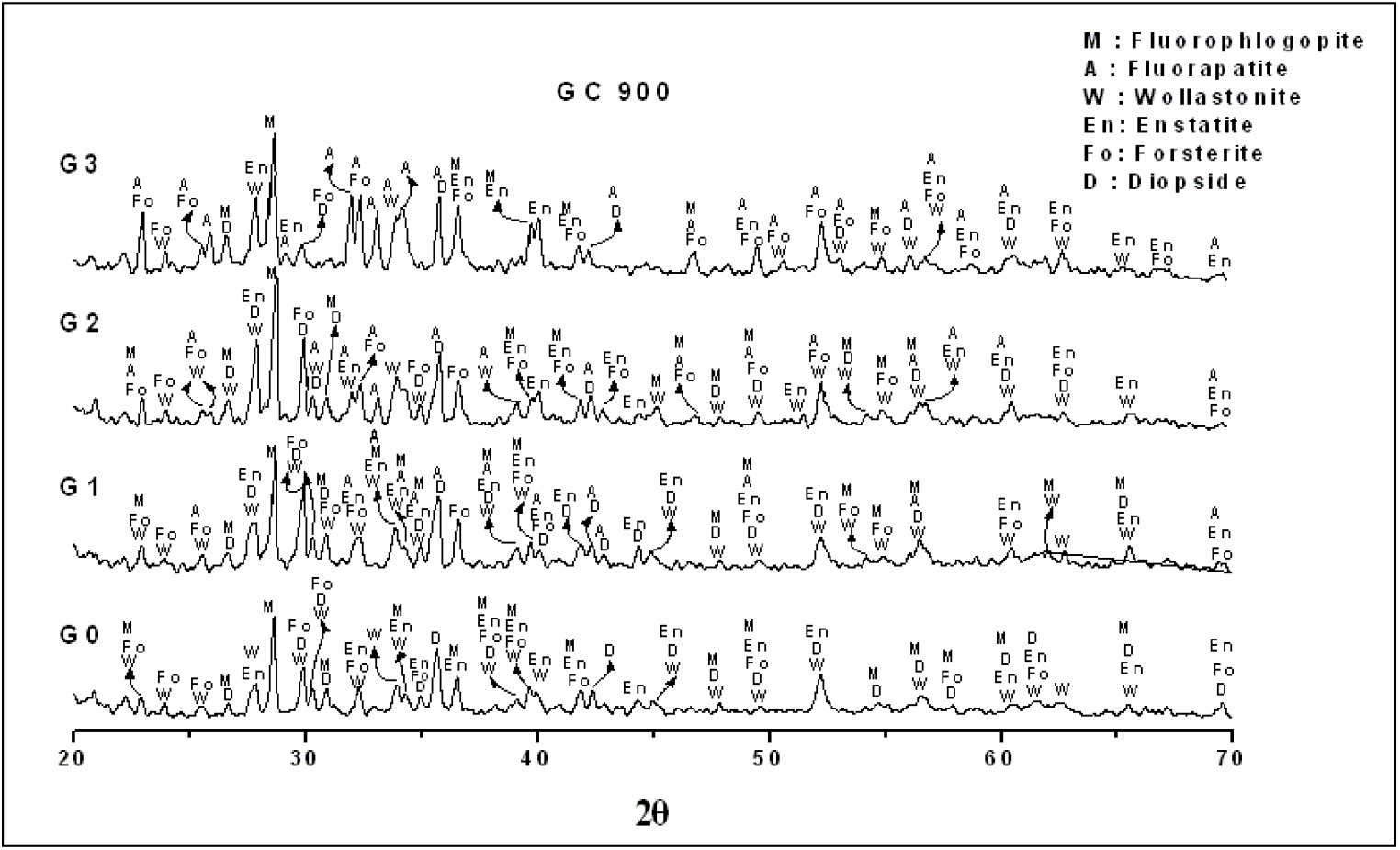
XRD patterns of the glass samples G0, G1, G2, and G3 crystallized at 900 °C for 2 h.

The addition of P_2_O_5_ significantly influenced mica crystallization, leading to a decrease in both crystal size and the degree of crystallinity of the mica phase, as shown in Tables S3 & S4. This effect can be attributed to P_2_O_5_ promoting liquid–liquid phase separation, which causes Ca and Mg ions to associate with phosphate-rich phases. As a result, the crystallization of the Si, Mg-rich mica phase was suppressed, leading to a reduction in its crystal size and crystallinity. Conversely, the crystallization of fluoroapatite (FA) increased with higher P_2_O_5_ content, as P_2_O_5_ interacts with Ca and F to form FA phases, leading to an increase in FA crystal size (Table S5). This effect was particularly evident at higher P_2_O_5_ concentrations (7.5 wt%).^12^

Additionally, Table S6 shows that the overall crystal size of the samples decreased with increasing P_2_O_5_ content. This may be attributed to P_2_O_5_ weakening the glass structure, reducing viscosity, and disrupting the rearrangement of the glass network, thereby inhibiting crystal growth.

Consequently, P_2_O_5_ increases the residual glass phase surrounding mica flakes by suppressing their crystallization. These observations are consistent with SEM microstructural analysis (Figure 5), which revealed an increase in residual glass phase surrounding mica flakes at elevated P_2_O_5_ levels.

**Figure 5.**
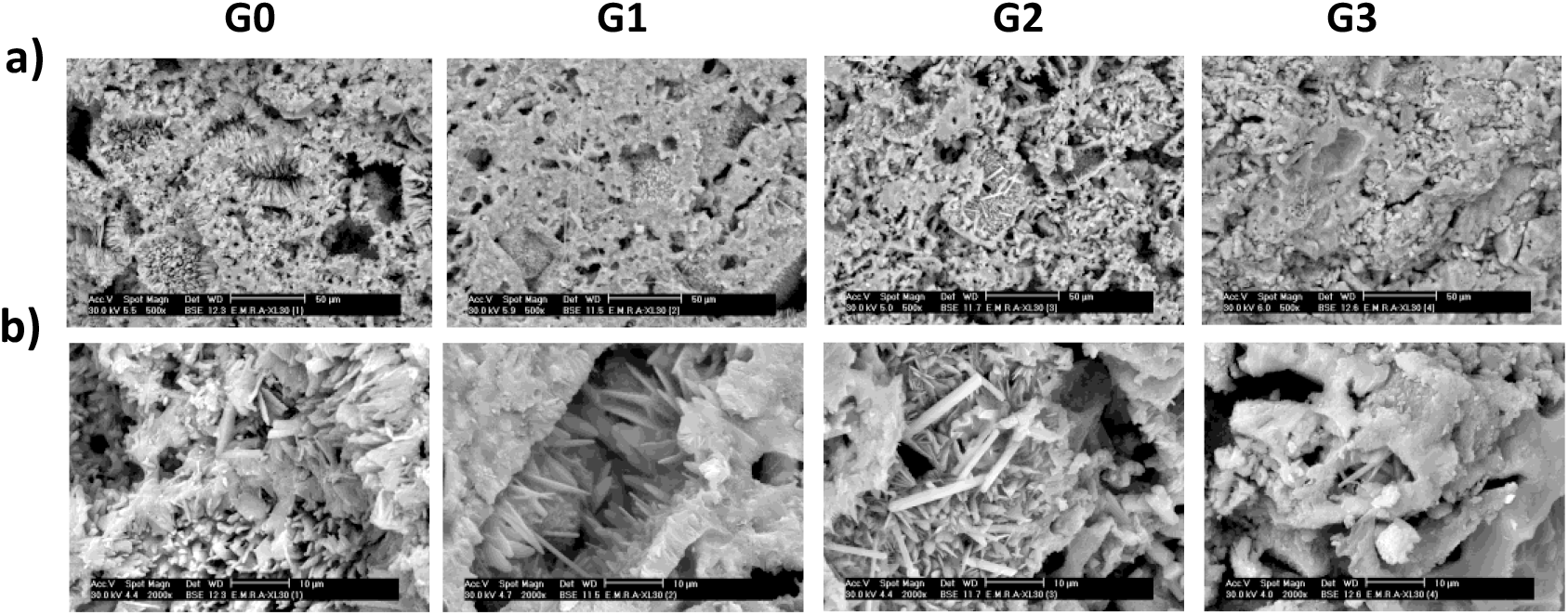
SEM of the investigated samples heat-treated at 900 °C for 2 h, (a) low magnification X500, and (b) high magnification X2000.

SEM analysis of samples heat-treated at 900°C for 2 hours revealed the influence of P_2_O_5_ on the parent glass composition (Figure 5). The G0 sample exhibits a dense, interlocked crystalline structure with well-defined fluorphlogopite flakes and embedded rod-like crystals, surrounded by some porosity. With the addition of P_2_O_5_ in G1, the residual glass phase increases, leading to more cemented fluormica flakes and the appearance of small rod-like crystals embedded in the glassy matrix. In G2, a further increase in P_2_O_5_ promotes the formation of a more homogeneous mixture of rod-like crystals with fewer distinct fluormica flakes. The G3 sample, with the highest P_2_O_5_ content, shows a predominant glassy phase with a significant reduction in fluormica flakes, indicating that P_2_O_5_ suppresses fluorphlogopite crystallization while promoting the growth of rod-like fluoroapatite crystals. These observations align with the XRD results, demonstrating the role of P_2_O_5_ in modulating the crystallization behavior of silicate-based glass-ceramics.

### 3.2 Physical and Mechanical Properties

The physical and mechanical properties of the glasses and glass-ceramics, including density, porosity, and microhardness, were evaluated to assess the influence of P_2_O_5_ on the structural integrity and performance of the materials. As shown in Figure 5, the theoretical density of the glassy samples is consistently higher than the experimental density for all compositions. This discrepancy can be attributed to the presence of micropores, which are not accounted for in the theoretical density calculations.

The experimental density of the glasses decreased with increasing P_2_O_5_ content, from 2.76 g/cm^3^ for G0 to 2.65 g/cm^3^ for G3 (Figure 6a). This reduction in density is attributed to the disruption of the Si-O-Si network by P_2_O_5_, which introduces more open structures and reduces the overall packing density of the glass.^11^ Additional comparisons between the densities of wax-coated and uncoated glass-ceramic samples are provided in the Supporting Information (Table S7). It can be observed that the density of uncoated glass-ceramic samples is consistently higher than that of coated samples. This difference is attributed to the presence of pores in the glass-ceramic structure, which decrease with increasing P_2_O_5_ content. These pores have the capacity to absorb the surrounding solution, leading to an increase in mass. As the number and size of the pores increase, more solution is retained within the structure, contributing to the higher density of the uncoated glass-ceramic samples compared to the coated ones.

**Figure 6.**
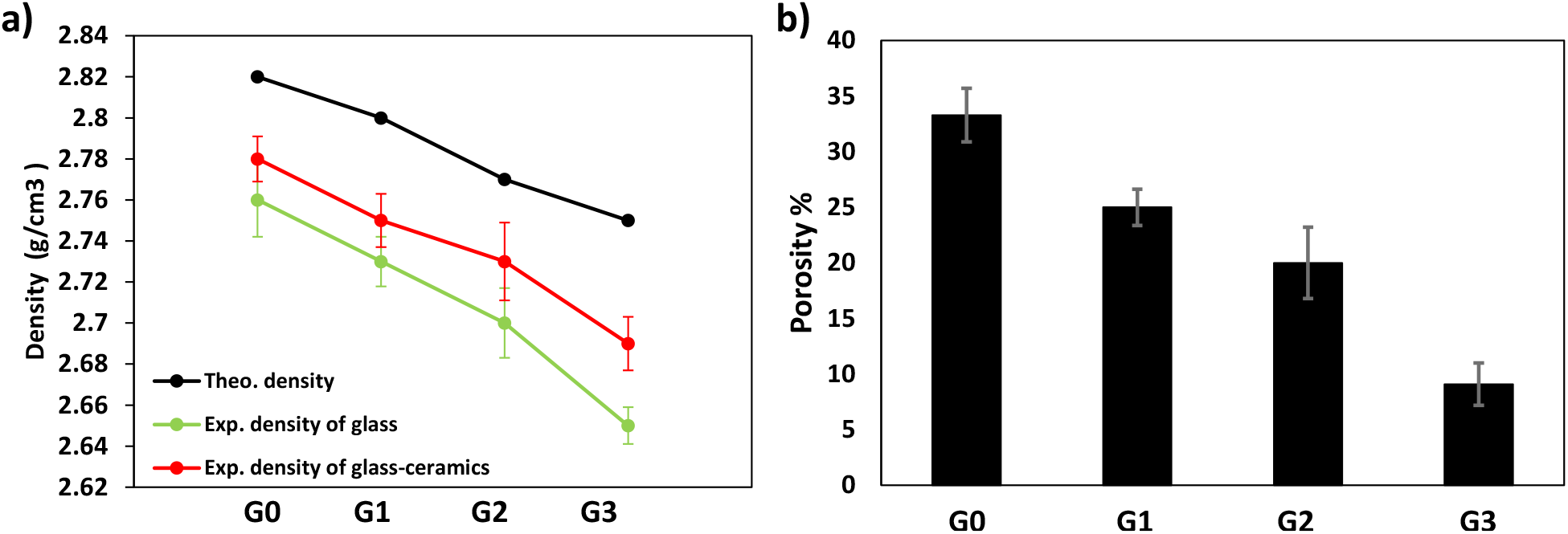
a) Comparison between the theoretical oxides and experimental density of glass and wax-coated glass-ceramic samples. b) The relationship between the glass-ceramics samples and their porosity (%).

Similarly, the porosity (Figure 6b) of the glass-ceramics also decreased with P_2_O_5_ addition, from 33.3% for G0 to 9.09% for G3. This decrease in porosity is likely due to the increased crystallization of fluoroapatite, which fills the voids and creates a more compact structure. The reduction in porosity is beneficial for the mechanical properties of the glass-ceramics, as it enhances their strength and fracture toughness.^10^

The microhardness of the glass-ceramics was measured to evaluate the effect of P_2_O_5_ on the mechanical properties. The microhardness value decreased from 380 kg/mm^2^ for G0 to 311 kg/mm^2^ for G3, as shown in Table 1. This decrease in microhardness is directly correlated with the increasing P_2_O_5_ content. The reduction in hardness can be attributed to the increased residual glass phase in the glass-ceramics with higher P_2_O_5_ content. P_2_O_5_ has a larger disrupting effect on the silicate network, weakening the overall structure by introducing non-bridging oxygen (NBO) atoms and reducing the connectivity of the glass matrix^12^. As a result, the glass-ceramics with higher P_2_O_5_ content exhibit lower mechanical strength and hardness due to the dominance of the residual glass phase, which is less rigid compared to the crystalline phases.

**Table 1.** Variation of microhardness with the glass ceramic samples’ compositions

| Sample Code | Vickers Microhardness (Kg/mm<sup>3</sup>) |
| --- | --- |
| **G0** | **380** |
| **G1** | **353** |
| **G2** | **327** |
| **G3** | **311** |

### 3.3 *In Vitro* Bioactivity

*In vitro* testing in simulated body fluid (SBF) revealed that P_2_O_5_ addition significantly enhances the bioactivity of glasses by optimizing HCA formation kinetics (Figures 7 & 8). As shown in Figure 7a, P_2_O_5_-free glass (G0) exhibited uncontrolled dissolution behavior, characterized by rapid pH elevation, indicating excessive alkali and alkaline earth ion (Na^+^, Ca^2+^, Mg^2+^) release from the glass surface, along with a rapid depletion of phosphate (P^5+^) in SBF (Figure 8a), suggesting non-productive P^5+^ consumption. In contrast, P_2_O_5_-containing glasses (G1-G3) demonstrated controlled bioactive behavior, showing moderated initial ion release (reduced pH spike) while maintaining P^5+^ availability during the critical 0-7 day period. This is evidenced by the distinct P^5+^ release peak observed in Figure 7. This controlled dissolution process follows established bioactive glass mechanisms: initial ion exchange is succeeded by network dissolution, leading to the release of silicic acid (Si(OH)4) into the solution and the formation of a silica-rich gel layer on the glass surface^6^. The subsequent precipitation of P^5+^ and Ca^2+^ ions from the solution leads to crystalline HCA formation, as confirmed by the consumption of P^5+^ and Ca^2+^ ions from the SBF solution after day 7 and day 15, respectively (Figure 7a).

**Figure 7.**
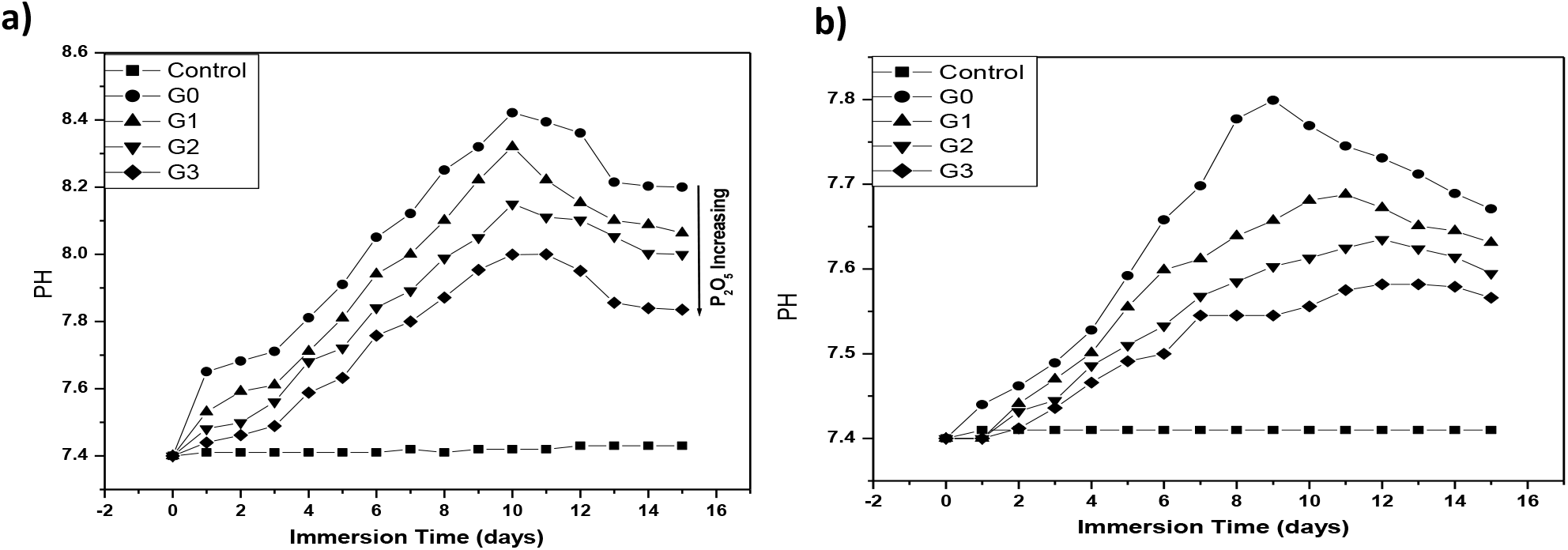
Shows pH variation of SBF for different samples of a) glass and b) glass-ceramics.

**Figure 8.**
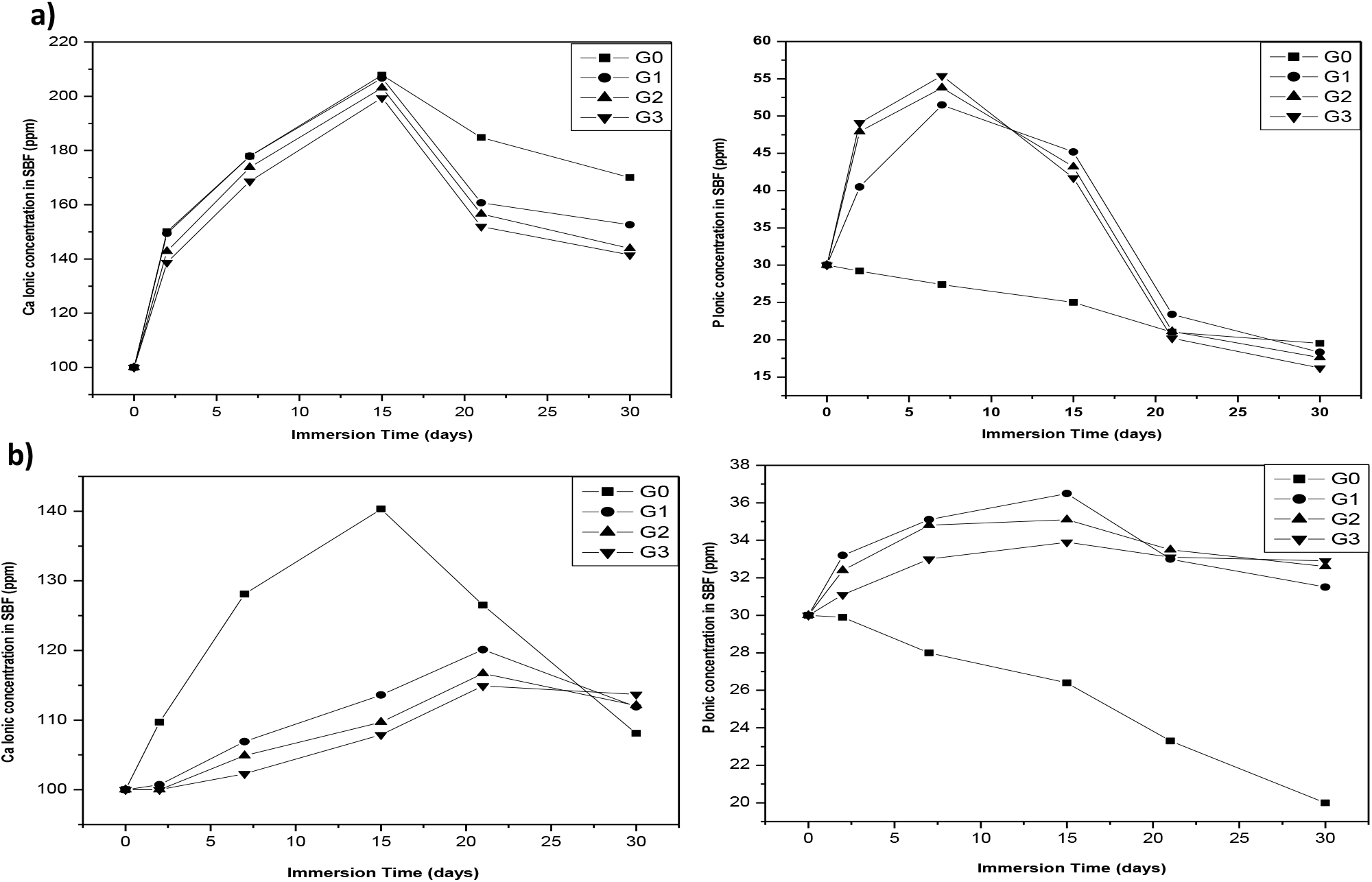
Ion concentration changes in (a) glass and (b) glass-ceramic samples after soaking in SBF for 30 days: Ca^2+^ (left) and P^5+^ (right).

However, in glass-ceramic compositions, the release of Ca^2+^ and P^5+^ ions was delayed due to the stabilization of the residual glass phase by P_2_O_5_, which suppresses ion exchange and network dissolution, as indicated by the slower pH increase in Figure 7b and the delayed release and subsequent consumption of Ca^2+^ and P^5+^ ions as shown in Figure 8b. As a result, HCA formation occurred at a reduced rate. The delayed bioactivity observed in glass-ceramics is attributed to increased crystallinity and the reduced reactivity of the residual glass phase. Despite this delay, bioactivity was still evident, as indicated by the formation of an HCA layer after 30 days of immersion in SBF. These findings suggest that while P_2_O_5_ enhances the bioactivity of bioactive glasses, its effect on glass-ceramics is more complex and is influenced by crystallization behavior and microstructural characteristics.^8^

FTIR spectroscopy was used to analyze the structural changes in the glasses and glass-ceramics before and after immersion in SBF. The spectra (Figure 9) of the unreacted glasses showed characteristic bands corresponding to Si-O-Si stretching vibrations (700-1175 cm^-1^) and Si-O-Si bending vibrations (415-540 cm^-1^). After immersion in SBF, new bands appeared at 470 cm^-1^, 560 cm^-1^, and 1060 cm^-1^, which are attributed to the formation of an HCA layer. The intensity of these bands increased with P_2_O_5_ content, indicating that P_2_O_5_ promotes the crystallization of HCA. In glass-ceramics (Figure 10), the intensity of the HCA bands was lower, suggesting that the increased crystallinity of the material due to the presence of P_2_O_5_ reduces its reactivity in SBF. The FTIR results are consistent with the XRD and SEM analyses and provide further evidence of the role of P_2_O_5_ in enhancing the bioactivity of silicate-based glasses.^7^ A detailed analysis of FTIR spectra is provided in the supporting information (Figures S9-S12 & Tables S9-S11).

**Figure 9.**
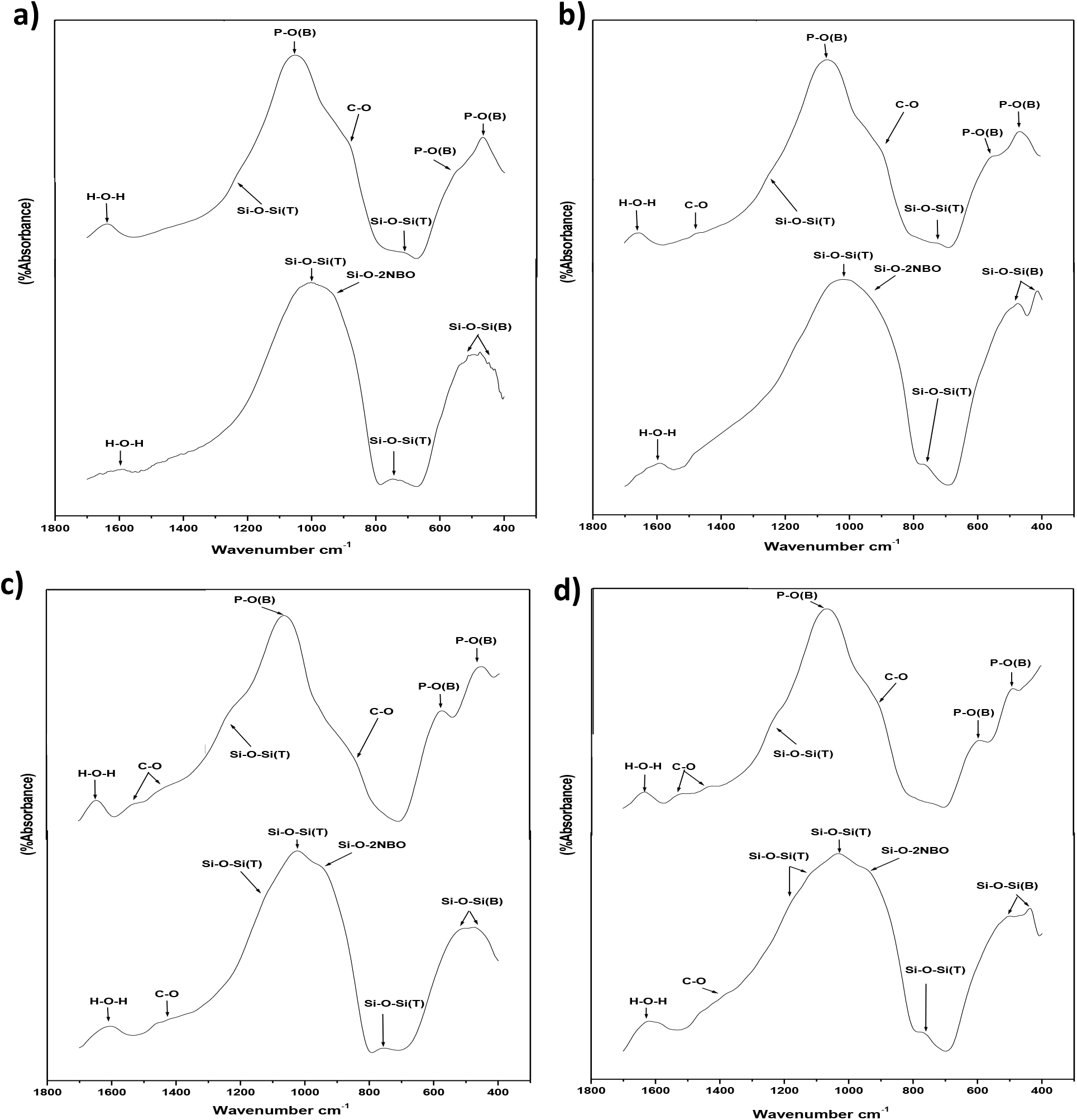
FTIR absorbance spectra for glass samples before (lower) and after (upper) soaking in SBF for 30 days: a) G0, b) G1, c) G2, and d) G3.

**Figure 10.**
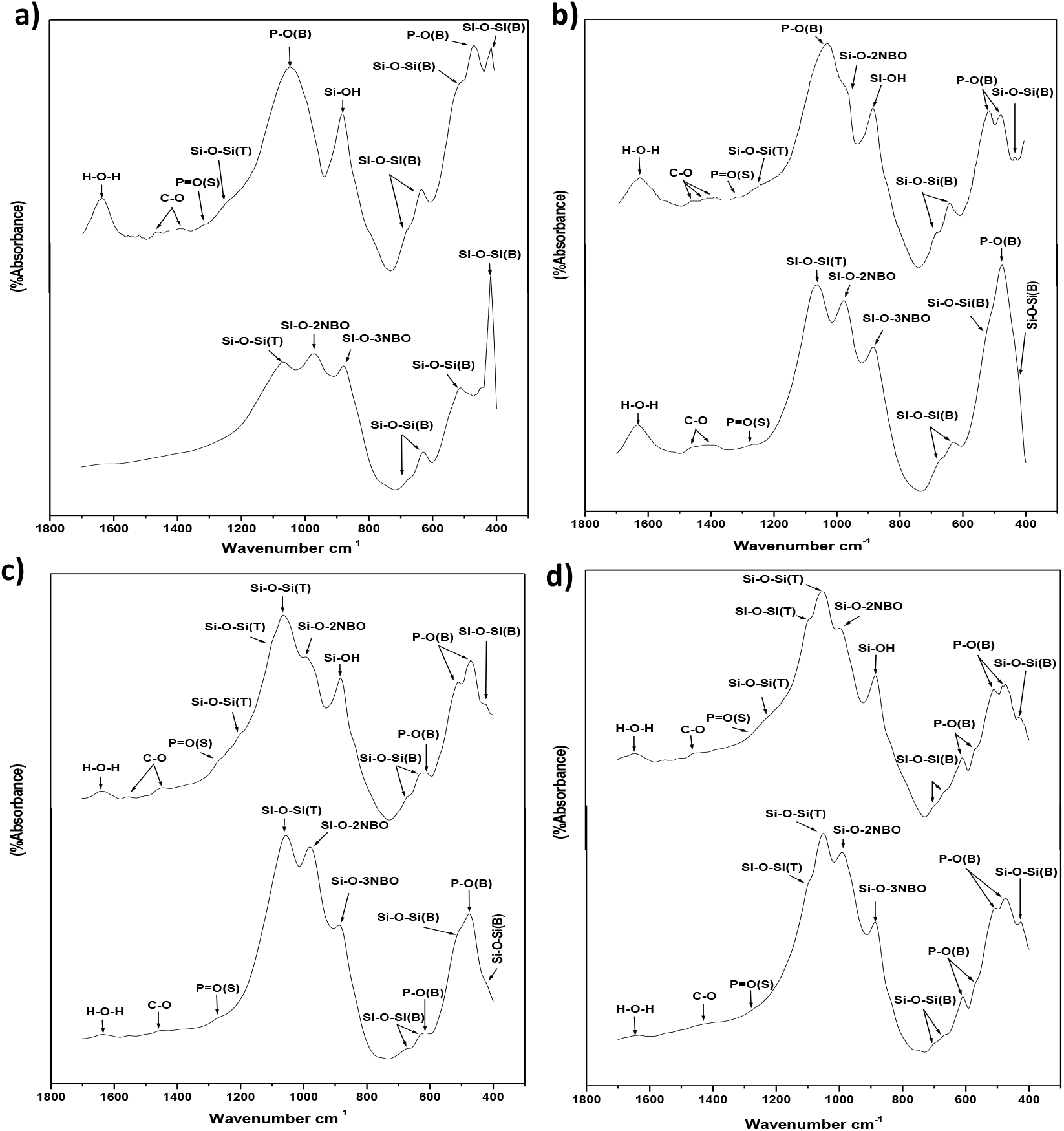
FTIR absorbance spectra for glass-ceramics samples before (lower) and after (upper) soaking in SBF for 30 days: a) G0, b) G1, c) G2, and d) G3.

## Conclusion

The addition of P_2_O_5_ to silicate-based bioactive glasses significantly influences their structure, crystallization behavior, and bioactivity. P_2_O_5_ lowers the glass transition temperature, promotes phase separation, and enhances the crystallization of fluoroapatite while reducing the formation of fluorophlogopite. In vitro tests demonstrated that P_2_O_5_ accelerates the formation of an HCA layer on the glass surface, indicating improved bioactivity. However, in glass-ceramics, P_2_O_5_ delays ion release and HCA formation due to the stabilization of the residual glass phase. These findings highlight the potential of P_2_O_5_ in tailoring the properties of bioactive glasses and glass-ceramics for specific biomedical applications.

## Acknowledgments

This research was conducted without external funding. We sincerely thank Prof. K. Tohamy and Prof. I. Hamzawi for their invaluable support and guidance throughout this study.

