## Supporting Information for "The Influence of P_₂_O_₅_ on the Structure, Crystallization, and Bioactivity of Silicate-Based Bioactive Glasses and Glass-Ceramics"

Mahmoud G. Soliman*, M. I. EL Gohary (deceased in 2022)

Biophysics Department, Faculty of Science (Boys), Al-Azhar University, Nasr City, Cairo, Egypt.

### Supporting Information

**Table S1**. Differential thermal analysis of G0, G1, G2 and G3 samples.

| | | | 1st peak (750-915) ºC | | | 2nd peak (750-950) ºC | | |
|---|---|---|---|---|---|---|---|---|
| **Sample code** | **Tg** | **Ts** | **Tc on** | **Tc max** | **Tc off** | **Tc on** | **Tc max** | **Tc off** |
| **G0** | 618 | 620 | 835 | 881 | 933 | - | - | - |
| **G1** | 612 | 620 | 777 | 828 | 880 | 890 | 915 | 978 |
| **G2** | 612 | 620 | 748 | 800 | 845 | 852 | 870 | 973 |
| **G3** | 595 | 612 | 700 | 750 | 785 | 890 | 915 | 990 |

**Table S2.** Shows the crystalline phases for G0, G1, G2, and G3 samples that are crystallized at 750 ºC and 900 ºC for 2 h

| Sample | **Heat- Treatment (ºC)** | **Products** | **Phases** |
|---|---|---|---|
| **G0** | 750 | Transparent glass | Am |
| | 900 | white | Fl, Fo, En, Dio, Wo |
| **G1** | 750 | white | Fl, FA, Dio |
| | 900 | white | Fl, FA, Fo, En, Dio, Wo |
| **G2** | 750 | white | Fl, FA, Fo, En, Dio, Wo |
| | 900 | white | Fl, FA, Fo, En, Dio, Wo |
| **G3** | 750 | white | Fl, FA, Fo, En, Dio, Wo |
| | 900 | white | Fl, FA, Fo, En, Dio, Wo |

**Am:** amorphous**, Fl.:** fluorophlogopite**, Fo:** forsterite**, En:** Enstatite**, Dio:** diopside**, Wo:** wollastonit

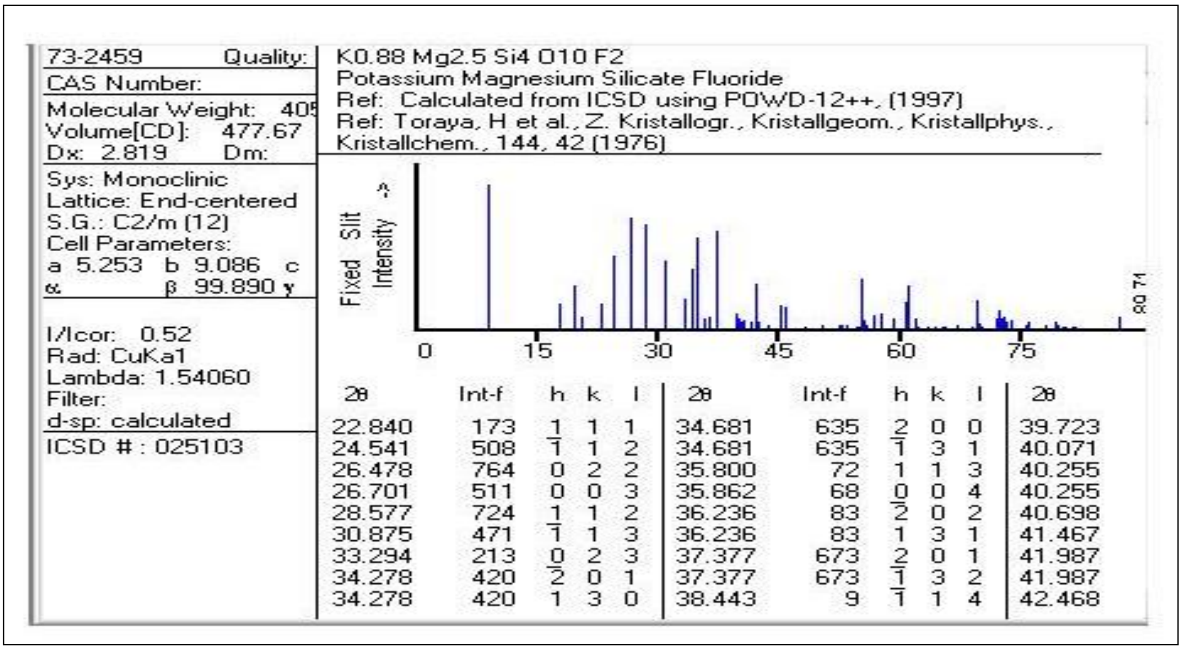

73-2459 Quality:
CAS Number:
Molecular Weight: 405
Volume[CD]: 477.67
Dx: 2.819 Dm:
Sys: Monoclinic
Lattice: End-centered
S.G.: C2/m (12)
Cell Parameters:
a 5.253 b 9.086 c
α β 99.890 γ
I/Icor: 0.52
Rad: CuKa1
Lambda: 1.54060
Filter:
d-sp: calculated
ICSD # : 025103

K0.88 Mg2.5 Si4 O10 F2
Potassium Magnesium Silicate Fluoride
Ref: Calculated from ICSD using POWD-12++, (1997)
Ref: Toraya, H et al., Z. Kristallogr., Kristallgeom., Kristallphys., Kristallchem., 144, 42 (1976)



| 2θ | Int-f | h | k | l | 2θ | Int-f | h | k | l | 2θ |
|---|---|---|---|---|---|---|---|---|---|---|
| 22.840 | 173 | 1 | 1 | 1 | 34.681 | 635 | 2 | 0 | 0 | 39.723 |
| 24.541 | 508 | $\bar{1}$ | 1 | 2 | 34.681 | 635 | $\bar{1}$ | 3 | 1 | 40.071 |
| 26.478 | 764 | 0 | 2 | 2 | 35.800 | 72 | 1 | 1 | 3 | 40.255 |
| 26.701 | 511 | 0 | 0 | 3 | 35.862 | 68 | 0 | 0 | 4 | 40.255 |
| 28.577 | 724 | 1 | 1 | 2 | 36.236 | 83 | $\bar{2}$ | 0 | 2 | 40.698 |
| 30.875 | 471 | $\bar{1}$ | 1 | 3 | 36.236 | 83 | 1 | 3 | 1 | 41.467 |
| 33.294 | 213 | 0 | 2 | 3 | 37.377 | 673 | 2 | 0 | 1 | 41.987 |
| 34.278 | 420 | $\bar{2}$ | 0 | 1 | 37.377 | 673 | $\bar{1}$ | 3 | 2 | 41.987 |
| 34.278 | 420 | 1 | 3 | 0 | 38.443 | 9 | $\bar{1}$ | 1 | 4 | 42.468 |

**Figure S1**. X-ray diffraction card No. (73-2459) corresponding to potassium magnesium silicate fluoride “M” ($K_{0.88}Mg_{2.5}Si_4O_{10}F_2$) phase formed upon heat treatment of G0, G1, G2, and G3 powder heat treated at 750 °C/2h and 900 °C/2h.

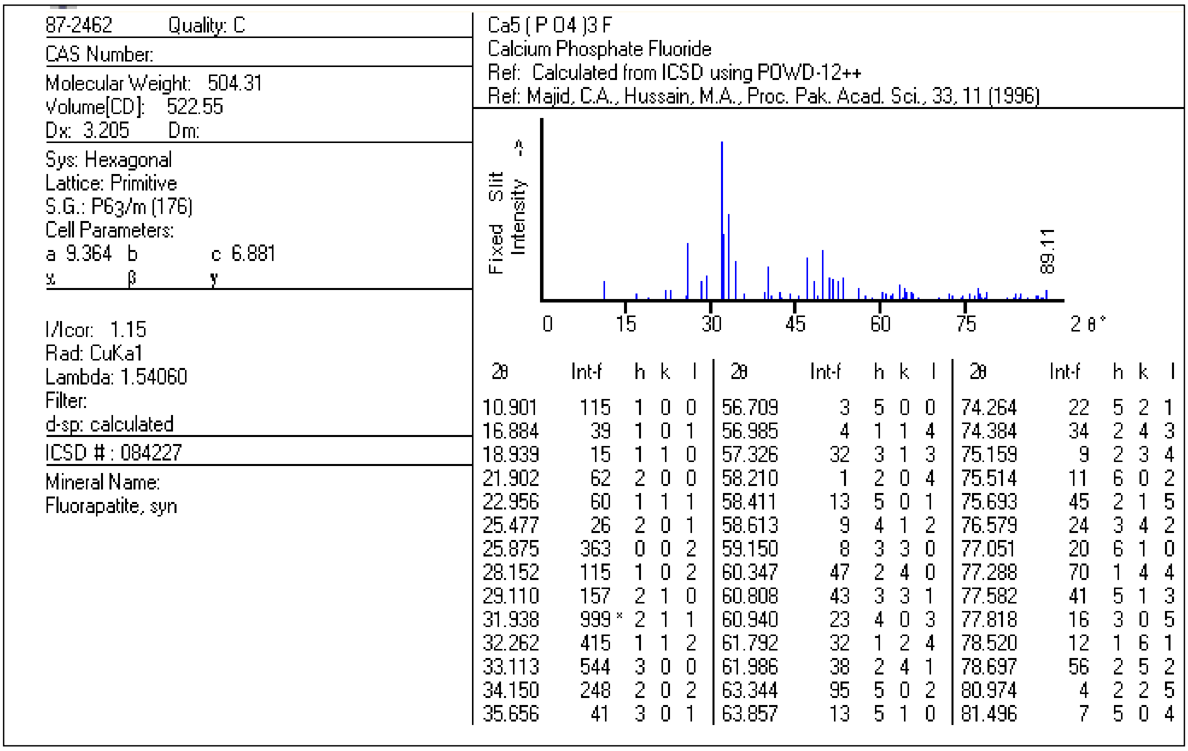

87-2462 Quality: C
CAS Number:
Molecular Weight: 504.31
Volume[CD]: 522.55
Dx: 3.205 Dm:
Sys: Hexagonal
Lattice: Primitive
S.G.: P63/m (176)
Cell Parameters:
a 9.364 b c 6.881
α β γ
I/Icor: 1.15
Rad: CuKa1
Lambda: 1.54060
Filter:
d-sp: calculated
ICSD # : 084227
Mineral Name:
Fluorapatite, syn

Ca5 ( P O4 )3 F
Calcium Phosphate Fluoride
Ref: Calculated from ICSD using POWD-12++
Ref: Majid, C.A., Hussain, M.A., Proc. Pak. Acad. Sci., 33, 11 (1996)



| 2θ | Int-f | h | k | l | 2θ | Int-f | h | k | l | 2θ | Int-f | h | k | l |
|---|---|---|---|---|---|---|---|---|---|---|---|---|---|---|
| 10.901 | 115 | 1 | 0 | 0 | 56.709 | 3 | 5 | 0 | 0 | 74.264 | 22 | 5 | 2 | 1 |
| 16.884 | 39 | 1 | 0 | 1 | 56.985 | 4 | 1 | 1 | 4 | 74.384 | 34 | 2 | 4 | 3 |
| 18.939 | 15 | 1 | 1 | 0 | 57.326 | 32 | 3 | 1 | 3 | 75.159 | 9 | 2 | 3 | 4 |
| 21.902 | 62 | 2 | 0 | 0 | 58.210 | 1 | 2 | 0 | 4 | 75.514 | 11 | 6 | 0 | 2 |
| 22.956 | 60 | 1 | 1 | 1 | 58.411 | 13 | 5 | 0 | 1 | 75.693 | 45 | 2 | 1 | 5 |
| 25.477 | 26 | 2 | 0 | 1 | 58.613 | 9 | 4 | 1 | 2 | 76.579 | 24 | 3 | 4 | 2 |
| 25.875 | 363 | 0 | 0 | 2 | 59.150 | 8 | 3 | 3 | 0 | 77.051 | 20 | 6 | 1 | 0 |
| 28.152 | 115 | 1 | 0 | 2 | 60.347 | 47 | 2 | 4 | 0 | 77.288 | 70 | 1 | 4 | 4 |
| 29.110 | 157 | 2 | 1 | 0 | 60.808 | 43 | 3 | 3 | 1 | 77.582 | 41 | 5 | 1 | 3 |
| 31.938 | 999 * | 2 | 1 | 1 | 60.940 | 23 | 4 | 0 | 3 | 77.818 | 16 | 3 | 0 | 5 |
| 32.262 | 415 | 1 | 1 | 2 | 61.792 | 32 | 1 | 2 | 4 | 78.520 | 12 | 1 | 6 | 1 |
| 33.113 | 544 | 3 | 0 | 0 | 61.986 | 38 | 2 | 4 | 1 | 78.697 | 56 | 2 | 5 | 2 |
| 34.150 | 248 | 2 | 0 | 2 | 63.344 | 95 | 5 | 0 | 2 | 80.974 | 4 | 2 | 2 | 5 |
| 35.656 | 41 | 3 | 0 | 1 | 63.857 | 13 | 5 | 1 | 0 | 81.496 | 7 | 5 | 0 | 4 |

**Figure S2**. X-ray diffraction card No. (87-2462) corresponding to fluorapatite “FA” ($Ca_5(PO_4)_3F$) phase formed upon heat treatment of G0, G1, G2 and G3 powder heat treated at 750 °C/2h and 900 °C/2h.

26-1322 Quality: I

CAS Number:

Molecular Weight: 406.21
Volume[CD]: 479.95
Dx: 2.811 Dm: 2.770

Sys: Monoclinic

S.G.:
Cell Parameters:
a 5.256 b 9.09 c 10.20
α β 99.98 γ

SS/FOM: F30=9(0.025,141)
I/Icor:
Rad: Cu
Lambda: 1.54056
Filter:
d-sp:

K Mg2.5 Si4 O10 ( O H )2
Potassium Magnesium Silicate Hydroxide
Ref: Seifert, Schreyer, Beitr. Mineral. Petrogr., 30, 196 (1971)

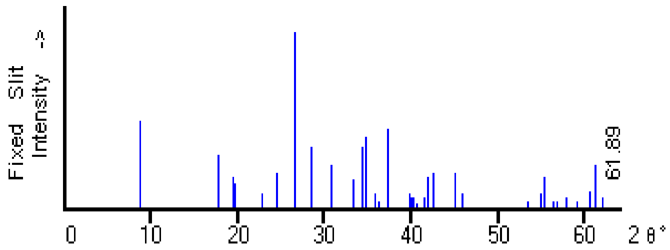


| 2θ | Int-f | h | k | l | 2θ | Int-f | h | k | l | 2θ | Int-f | h | k | l |
|---|---|---|---|---|---|---|---|---|---|---|---|---|---|---|
| 8.748 | 50 | 0 | 0 | 1 | 34.631 | 40 | 2 | 0 | 0 | 45.862 | 8 | 1 | 3 | 3 |
| 17.653 | 30 | 0 | 0 | 2 | 35.714 | 8 | 0 | 0 | 4 | 53.377 | 4 | 1 | 5 | 0 |
| 19.493 | 18 | 0 | 2 | 0 | 36.190 | 4 | 1 | 3 | 1 | 54.864 | 8 | 0 | 0 | 6 |
| 19.712 | 14 | 1 | 1 | 0 | 37.328 | 45 | $\bar{1}$ | 3 | 2 | 55.186 | 18 | 2 | 0 | 4 |
| 22.800 | 8 | 1 | 1 | 1 | 39.654 | 8 | 0 | 4 | 0 | 56.289 | 4 | $\bar{3}$ | 2 | 1 |
| 24.482 | 20 | $\bar{1}$ | 1 | 2 | 40.058 | 6 | 2 | 2 | 0 | 56.667 | 4 | $\bar{2}$ | 2 | 5 |
| 26.530 | 80 | $\bar{1}$ | 2 | 1 | 40.170 | 6 | $\bar{2}$ | 0 | 3 | 57.754 | 6 | 1 | 5 | 2 |
| 26.603 | 100b | 0 | 0 | 3 | 40.662 | 3 | 0 | 4 | 1 | 59.011 | 4 | $\bar{1}$ | 5 | 3 |
| 28.503 | 35 | 1 | 2 | 1 | 41.443 | 6 | $\bar{2}$ | 2 | 2 | 59.094 | 4 | 2 | 2 | 4 |
| 30.796 | 25 | 0 | 3 | 1 | 41.906 | 18 | 2 | 0 | 2 | 60.457 | 10 | 1 | 3 | 5 |
| 33.190 | 16 | 0 | 2 | 3 | 42.548 | 20 | 1 | 0 | 4 | 61.119 | 25 | 0 | 6 | 0 |
| 34.236 | 35 | $\bar{2}$ | 0 | 1 | 45.091 | 20 | 0 | 0 | 5 | 61.889 | 6 | 0 | 6 | 1 |

**Figure S3**. X-ray diffraction card No. (26-1322) corresponding to potassium magnesium silicate hydroxide "M" ($KMg_{2.5}Si_4O_{10}(OH)_2$) phase formed upon heat treatment of G0, G1, G2 and G3 powder heat treated at 750 °C/2h and 900 °C/2h.

7-0700 Quality: C
AS Number:
olecular Weight: 224.75
olume[CD]: 442.44
x: 3.374 Dm:
ys: Monoclinic
attice: End-centered
.G.: C2/c (15)
ell Parameters:
9.773 b 8.952 c 5.252
β 105.67 γ

/Icor: 1.33
ad: CuKa1
ambda: 1.54060
ilter:
-sp: calculated
CSD # : 064977
ineral Name:
iopside, syn
on-Ambient Temperature

Ca Mg.74 Fe.26 ( Si2 O6 )
Calcium Magnesium Iron Silicate
Ref: Calculated from ICSD using POWD-12++, (1997)
Ref: Raudsepp, M., Hawthorne, F.C., Turnock, A.C., Can. Mineral., 28, 93 (1990)

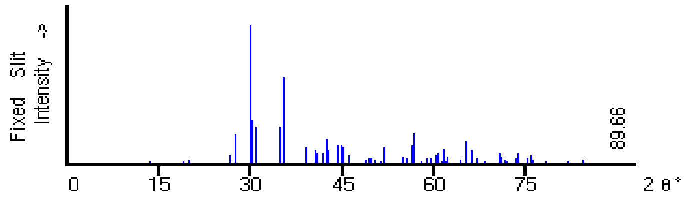


| 2θ | Int-f | h | k | l | 2θ | Int-f | h | k | l | 2θ | Int-f | h | k | l |
|---|---|---|---|---|---|---|---|---|---|---|---|---|---|---|
| 13.642 | 12 | 1 | 1 | 0 | 49.637 | 37 | 2 | 2 | 2 | 64.256 | 23 | $\bar{5}$ | 1 | 3 |
| 18.847 | 17 | 2 | 0 | 0 | 50.152 | 26 | $\bar{3}$ | 3 | 2 | 64.256 | 23 | $\bar{6}$ | 2 | 2 |
| 19.819 | 34 | 0 | 2 | 0 | 50.152 | 26 | 1 | 3 | 2 | 64.350 | 15 | 4 | 2 | 2 |
| 20.139 | 3 | $\bar{1}$ | 1 | 1 | 50.498 | 8 | 2 | 4 | 1 | 65.132 | 26 | 0 | 6 | 1 |
| 24.230 | 11 | 1 | 1 | 1 | 51.244 | 15 | 4 | 2 | 1 | 65.301 | 161 | 5 | 3 | 1 |
| 26.573 | 65 | 0 | 2 | 1 | 51.946 | 124 | 1 | 5 | 0 | 66.145 | 99 | $\bar{3}$ | 5 | 2 |
| 27.482 | 206 | 2 | 2 | 0 | 53.183 | 11 | $\bar{5}$ | 1 | 2 | 66.145 | 99 | 1 | 5 | 2 |
| 29.807 | 999 * | $\bar{2}$ | 2 | 1 | 53.267 | 11 | 3 | 1 | 2 | 66.718 | 6 | 1 | 3 | 3 |
| 30.167 | 316 | 3 | 1 | 0 | 54.304 | 11 | $\bar{1}$ | 5 | 1 | 67.129 | 36 | 2 | 2 | 3 |
| 30.842 | 270 | $\bar{3}$ | 1 | 1 | 54.728 | 49 | $\bar{2}$ | 4 | 2 | 67.892 | 8 | $\bar{7}$ | 1 | 1 |
| 31.424 | 9 | 1 | 3 | 0 | 54.728 | 49 | 0 | 4 | 2 | 68.118 | 13 | $\bar{2}$ | 4 | 3 |
| 34.875 | 266 | $\bar{1}$ | 3 | 1 | 55.374 | 40 | $\bar{3}$ | 1 | 3 | 69.791 | 10 | 0 | 4 | 3 |
| 35.531 | 622 | 0 | 0 | 2 | 56.184 | 8 | 1 | 5 | 1 | 70.126 | 4 | 2 | 6 | 1 |
| 35.531 | 622 | 2 | 2 | 1 | 56.184 | 8 | $\bar{4}$ | 4 | 1 | 70.641 | 77 | $\bar{7}$ | 1 | 2 |
| 37.491 | 3 | 1 | 3 | 1 | 56.452 | 134 | $\bar{5}$ | 3 | 1 | 70.641 | 77 | 6 | 2 | 1 |
| 38.229 | 10 | 4 | 0 | 0 | 56.607 | 219 | $\bar{2}$ | 2 | 3 | 70.827 | 51 | 7 | 1 | 0 |
| 38.947 | 122 | 3 | 1 | 1 | 56.727 | 114 | 4 | 4 | 0 | 70.996 | 16 | 3 | 1 | 3 |

**Figure S4**. X-ray diffraction card No. (87-0700) corresponding to calcium magnesium iron silicate "D" ($CaMg_{.74}Fe_{.26}(Si_2O_6)$) phase formed upon heat treatment of G0, G1, G2, and G3 powder heat treated at 750 °C/2h and 900 °C/2h.

87-0704 Quality: C
CAS Number:
Molecular Weight: 100.39
Volume[CD]: 381.50
Dx: 3.496 Dm:
Sys: Monoclinic
Lattice: End-centered
S.G.: C2/c (15)
Cell Parameters:
a 9.201 b 8.621 c 4.908
α β 101.50 γ
I/Icor: 0.56
Rad: CuKa1
Lambda: 1.54060
Filter:
d-sp: calculated
ICSD # : 064981
Mineral Name:
Clinoenstatite HP
Non-Ambient Pressure

Mg ( Si O3 )
Magnesium Silicate
Ref: Calculated from ICSD using POWD-12++, (1997)
Ref: Angel, R.J., Chopelas, A., Ross, N.L., Nature (London), 358, 322 (1992)

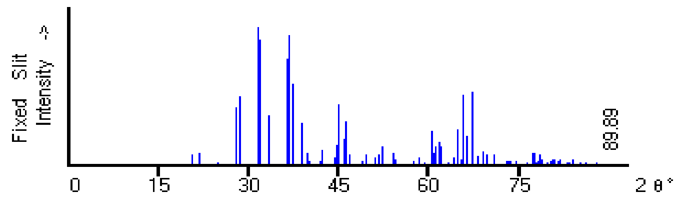


| 2θ | Int-f | h | k | l | 2θ | Int-f | h | k | l | 2θ | Int-f | h | k | l |
|---|---|---|---|---|---|---|---|---|---|---|---|---|---|---|
| 14.203 | 3 | 1 | 1 | 0 | 51.513 | 7 | $\bar{5}$ | 1 | 1 | 68.951 | 93 | 1 | 5 | 2 |
| 19.677 | 1 | 2 | 0 | 0 | 51.751 | 78 | 5 | 1 | 0 | 69.481 | 70 | 2 | 2 | 3 |
| 20.588 | 79 | 0 | 2 | 0 | 52.193 | 136 | 2 | 4 | 1 | 69.683 | 49 | 1 | 3 | 3 |
| 21.714 | 84 | $\bar{1}$ | 1 | 1 | 52.583 | 5 | 4 | 2 | 1 | 70.287 | 6 | $\bar{2}$ | 6 | 1 |
| 24.876 | 16 | 1 | 1 | 1 | 54.111 | 86 | 1 | 5 | 0 | 70.287 | 6 | $\bar{6}$ | 2 | 2 |
| 27.770 | 413 | 0 | 2 | 1 | 54.222 | 48 | $\bar{4}$ | 2 | 2 | 70.750 | 70 | $\bar{3}$ | 5 | 2 |
| 28.629 | 494 | 2 | 2 | 0 | 54.411 | 36 | 3 | 1 | 2 | 71.275 | 8 | $\bar{5}$ | 1 | 3 |
| 31.499 | 999 * | 3 | 1 | 0 | 56.919 | 1 | $\bar{1}$ | 5 | 1 | 72.257 | 7 | 5 | 1 | 2 |
| 31.977 | 905 | $\bar{2}$ | 2 | 1 | 57.364 | 34 | 0 | 4 | 2 | 72.913 | 34 | 6 | 2 | 1 |
| 32.680 | 6 | 1 | 3 | 0 | 58.467 | 53 | $\bar{5}$ | 1 | 2 | 72.913 | 34 | $\bar{7}$ | 1 | 1 |
| 33.480 | 356 | $\bar{3}$ | 1 | 1 | 59.122 | 10 | 5 | 1 | 1 | 73.150 | 26 | $\bar{2}$ | 4 | 3 |
| 36.427 | 765 | 2 | 2 | 1 | 59.273 | 15 | 4 | 4 | 0 | 73.574 | 31 | 0 | 4 | 3 |
| 36.745 | 937 | $\bar{1}$ | 3 | 1 | 59.782 | 3 | $\bar{4}$ | 4 | 1 | 74.411 | 27 | 7 | 1 | 0 |
| 37.365 | 590 | 0 | 0 | 2 | 60.357 | 240 | $\bar{5}$ | 3 | 1 | 76.165 | 5 | $\bar{5}$ | 5 | 1 |
| 38.183 | 5 | $\bar{1}$ | 1 | 2 | 60.571 | 133 | 5 | 3 | 0 | 76.357 | 19 | 5 | 5 | 0 |
| 38.736 | 301 | $\bar{2}$ | 0 | 2 | 60.717 | 80 | $\bar{3}$ | 1 | 3 | 76.517 | 10 | $\bar{6}$ | 4 | 1 |

**Figure S5**. X-ray diffraction card No. (87-0704) corresponding to magnesium silicate "En" (Mg $(SiO_3)$) phase formed upon heat treatment of G0, G1, G2, and G3 powder heat treated at 750 **°C/2h and 900 °C/2h.**

84-0655 Quality: C
CAS Number:
Molecular Weight: 116.16
Volume[CD]: 795.07
Dx: 2.911 Dm:
Sys: Monoclinic
Lattice: Primitive
S.G.: P21/a (14)
Cell Parameters:
a ¯5.42 b 7.324 c 7.069
α β 95.371 γ
I/Icor: 1.81
Rad: CuKa1
Lambda: 1.54060
Filter:
d-sp: calculated
ICSD # : 201538
Mineral Name:
Wollastonite 2M

Ca Si O3
Calcium Silicate
Ref: Calculated from ICSD using POWD-12++, (1997)
Ref: Ohashi, Y., Phys. Chem. Miner., 10, 217 (1984)

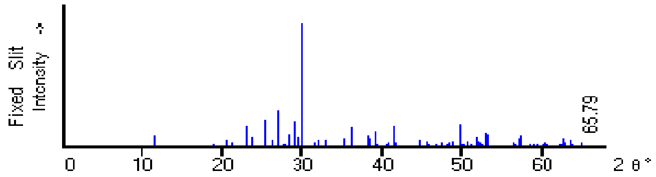


| 2θ | Int-f | h | k | l | 2θ | Int-f | h | k | l | 2θ | Int-f | h | k | l |
|---|---|---|---|---|---|---|---|---|---|---|---|---|---|---|
| 25.288 | 214 | 0 | 0 | 2 | 39.050 | 117 | 2 | 0 | $\bar{3}$ | 47.413 | 18 | 6 | 1 | 2 |
| 25.288 | 214 | 4 | 0 | $\bar{1}$ | 39.465 | 13 | 5 | 2 | $\bar{1}$ | 47.773 | 7 | 3 | 3 | $\bar{2}$ |
| 26.187 | 46 | 4 | ¯ | 0 | 40.235 | 5 | 1 | 1 | $\bar{3}$ | 47.986 | 16 | 5 | 2 | 2 |
| 26.841 | 284 | 2 | 0 | $\bar{2}$ | 40.336 | 8 | 0 | 1 | 3 | 47.986 | 16 | 3 | 2 | $\bar{3}$ |
| 27.464 | 18 | 0 | 2 | 1 | 40.499 | 12 | 2 | 3 | $\bar{1}$ | 48.142 | 11 | 5 | 1 | $\bar{3}$ |
| 27.464 | 18 | 4 | 0 | 1 | 40.499 | 12 | 6 | 1 | 1 | 48.324 | 45 | 7 | 2 | 0 |
| 27.801 | 21 | 1 | 2 | $\bar{1}$ | 40.707 | 34 | 3 | 2 | 2 | 48.324 | 45 | 7 | 1 | $\bar{2}$ |
| 28.221 | 93 | 4 | ¯ | $\bar{1}$ | 41.283 | 167 | 5 | 2 | 1 | 48.433 | 26 | 2 | 2 | 3 |
| 28.221 | 93 | 1 | ¯ | $\bar{2}$ | 41.283 | 167 | 1 | 1 | 3 | 48.593 | 14 | 5 | 3 | $\bar{1}$ |
| 28.858 | 204 | 2 | 0 | 2 | 41.611 | 29 | 6 | 0 | $\bar{2}$ | 48.763 | 8 | 6 | 2 | $\bar{2}$ |
| 29.525 | 71 | 2 | ¯ | $\bar{2}$ | 42.455 | 1 | 3 | 3 | $\bar{1}$ | 48.922 | 12 | 4 | 1 | 3 |
| 29.978 | 999 * | 3 | 2 | 0 | 42.653 | 5 | 5 | 1 | 2 | 49.054 | 43 | 8 | 1 | 0 |
| 31.384 | 6 | 2 | ¯ | 2 | 42.653 | 5 | 3 | 1 | $\bar{3}$ | 49.054 | 43 | 7 | 2 | $\bar{1}$ |
| 31.563 | 27 | 5 | ¯ | 0 | 43.236 | 8 | 4 | 0 | $\bar{3}$ | 49.757 | 171 | 0 | 4 | 0 |
| 31.959 | 55 | 3 | 2 | $\bar{1}$ | 43.492 | 1 | 6 | 1 | $\bar{2}$ | 50.135 | 22 | 4 | 3 | $\bar{2}$ |
| 32.849 | 50 | 4 | 0 | $\bar{2}$ | 43.492 | 1 | 3 | 3 | 1 | 50.135 | 22 | 5 | 3 | 1 |
| 33.270 | 2 | 3 | 2 | 1 | 43.803 | 1 | 7 | 1 | $\bar{1}$ | 50.393 | 17 | 8 | 0 | 1 |
| 33.800 | 1 | 4 | 2 | 0 | 43.915 | 1 | 4 | 3 | 0 | 50.795 | 43 | 3 | 2 | 3 |
| 34.470 | 3 | 3 | ¯ | 2 | 44.051 | 1 | 6 | 2 | $\bar{1}$ | 51.202 | 14 | 7 | 2 | 1 |
| 35.032 | 57 | 6 | 0 | 0 | 44.051 | 1 | 4 | 2 | 2 | 51.202 | 14 | 2 | 4 | 0 |

**Figure S6**. X-ray diffraction card No. (87-0655) corresponding to calcium silicate "W" (Ca $SiO_3$) phase formed upon heat treatment of G0, G1, G2, and G3 powder heat treated at 750 °C/2h and 900 °C/2h.

85-1364 Quality: C

CAS Number:

Molecular Weight: 140.69
Volume[CD]: 289.39
Dx: 3.229 Dm:

Sys: Orthorhombic
Lattice: Primitive
S.G.: Pbnm (62)
Cell Parameters:
a 4.749 b 10.19 c 5.980
α β γ

I/Icor: 0.75
Rad: CuKa1
Lambda: 1.54060
Filter:
d-sp: calculated

ICSD # : 064744

Mineral Name:
Forsterite, syn

Mg2 Si O4
Magnesium Silicate
Ref: Calculated from ICSD using POWD-12++, (1997)
Ref: Hazen, R.M., Am. Mineral., 61, 1280 (1976)

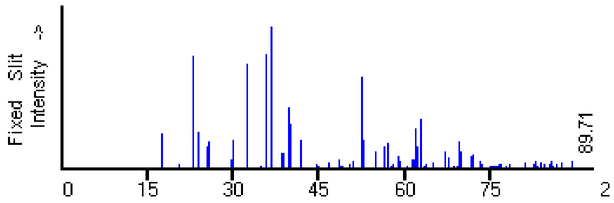


| 2θ | Int-f | h | k | l | 2θ | Int-f | h | k | l | 2θ | Int-f | h | k | l |
|---|---|---|---|---|---|---|---|---|---|---|---|---|---|---|
| 23.908 | 259 | 1 | 0 | 1 | 55.007 | 117 | 2 | 4 | 1 | 71.918 | 91 | 3 | 4 | 1 |
| 25.476 | 150 | 1 | 1 | 1 | 56.263 | 152 | 0 | 6 | 1 | 72.062 | 32 | 2 | 4 | 3 |
| 25.622 | 190 | 1 | 2 | 0 | 56.363 | 95 | 2 | 3 | 2 | 73.148 | 47 | 3 | 3 | 2 |
| 29.718 | 67 | 1 | 2 | 1 | 56.917 | 174 | 1 | 3 | 3 | 73.148 | 47 | 0 | 6 | 3 |
| 29.858 | 202 | 0 | 0 | 2 | 57.592 | 18 | 1 | 6 | 0 | 73.371 | 25 | 0 | 4 | 4 |
| 32.379 | 727 | 1 | 3 | 0 | 58.068 | 30 | 1 | 5 | 2 | 74.422 | 3 | 0 | 8 | 0 |
| 34.761 | 14 | 0 | 2 | 2 | 58.770 | 82 | 0 | 4 | 3 | 75.016 | 16 | 2 | 0 | 4 |
| 35.201 | 6 | 0 | 4 | 0 | 59.003 | 51 | 3 | 1 | 0 | 75.249 | 12 | 1 | 7 | 2 |
| 35.773 | 798 | 1 | 3 | 1 | 59.748 | 5 | 2 | 5 | 0 | 75.693 | 17 | 2 | 1 | 4 |
| 36.562 | 999 * | 1 | 1 | 2 | 60.446 | 18 | 3 | 0 | 1 | 75.798 | 13 | 2 | 6 | 2 |
| 37.859 | 7 | 2 | 0 | 0 | 61.196 | 38 | 3 | 1 | 1 | 76.072 | 10 | 3 | 5 | 0 |
| 38.376 | 107 | 0 | 4 | 1 | 61.268 | 42 | 3 | 2 | 0 | 76.274 | 15 | 1 | 6 | 3 |
| 38.914 | 111 | 2 | 1 | 0 | 61.352 | 58 | 2 | 1 | 3 | 76.494 | 27 | 1 | 4 | 4 |
| 39.743 | 425 | 1 | 2 | 2 | 61.711 | 62 | 2 | 4 | 2 | 76.732 | 22 | 2 | 7 | 0 |
| 40.136 | 311 | 1 | 4 | 0 | 62.028 | 278 | 0 | 0 | 4 | 76.835 | 28 | 3 | 0 | 3 |
| 41.849 | 201 | 2 | 1 | 1 | 62.235 | 149 | 1 | 4 | 3 | 77.507 | 7 | 3 | 1 | 3 |

**Figure S7**. X-ray diffraction card No. (85-1364) corresponding to magnesium silicate "Fo" ($Mg_2SiO_4$) phase formed upon heat treatment of G0, G1, G2, and G3 powder heat treated at 750 °C/2h and 900 °C/2h.

82-1406 Quality: C

CAS Number:

Molecular Weight: 60.08
Volume[CD]: 165.20
Dx: 2.416 Dm:

Sys: Tetragonal
Lattice: Primitive
S.G.: P41212 (92)
Cell Parameters:
a 4.922 b c 6.817
α β γ

I/Icor: 4.64
Rad: CuKa1
Lambda: 1.54060
Filter:
d-sp: calculated

ICSD # : 075486

Mineral Name:
Cristobalite alpha, syn

Si O2
Silicon Oxide
Ref: Calculated from ICSD using POWD-12++, (1997)
Ref: Parise, J.B et al., J. Appl. Phys., 75, 1361 (1994)

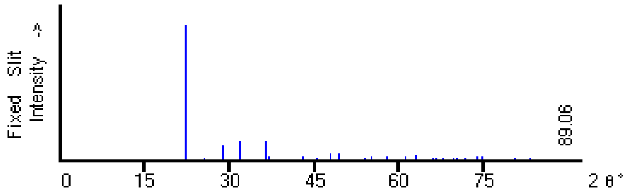


| 2θ | Int-f | h | k | l | 2θ | Int-f | h | k | l | 2θ | Int-f | h | k | l |
|---|---|---|---|---|---|---|---|---|---|---|---|---|---|---|
| 22.257 | 999 * | 1 | 0 | 1 | 57.148 | 4 | 1 | 0 | 4 | 74.858 | 26 | 3 | 2 | 2 |
| 25.571 | 12 | 1 | 1 | 0 | 57.744 | 30 | 3 | 0 | 1 | 78.483 | 5 | 2 | 2 | 4 |
| 28.775 | 107 | 1 | 1 | 1 | 58.310 | 7 | 2 | 1 | 3 | 78.995 | 10 | 4 | 0 | 1 |
| 31.909 | 137 | 1 | 0 | 2 | 59.319 | 7 | 3 | 1 | 0 | 80.360 | 12 | 4 | 1 | 0 |
| 36.476 | 145 | 2 | 0 | 0 | 59.597 | 5 | 2 | 2 | 2 | 81.333 | 1 | 3 | 0 | 4 |
| 36.878 | 26 | 1 | 1 | 2 | 60.429 | 1 | 1 | 1 | 4 | 81.841 | 6 | 4 | 1 | 1 |
| 38.870 | 1 | 2 | 0 | 1 | 61.005 | 31 | 3 | 1 | 1 | 82.326 | 10 | 3 | 2 | 3 |
| 40.963 | 1 | 2 | 1 | 0 | 62.800 | 41 | 3 | 0 | 2 | 83.294 | 15 | 2 | 1 | 5 |
| 43.147 | 32 | 2 | 1 | 1 | 65.913 | 13 | 3 | 1 | 2 | 83.438 | 7 | 4 | 0 | 2 |
| 43.845 | 2 | 1 | 0 | 3 | 66.699 | 15 | 2 | 0 | 4 | 84.162 | 9 | 3 | 1 | 4 |
| 45.415 | 15 | 2 | 0 | 2 | 67.765 | 13 | 2 | 2 | 3 | 84.667 | 4 | 3 | 3 | 1 |
| 47.759 | 55 | 1 | 1 | 3 | 68.694 | 1 | 3 | 2 | 0 | 86.255 | 1 | 4 | 1 | 2 |
| 49.232 | 52 | 2 | 1 | 2 | 69.721 | 22 | 2 | 1 | 4 | 88.178 | 1 | 1 | 0 | 6 |
| 52.540 | 6 | 2 | 2 | 0 | 70.257 | 13 | 3 | 2 | 1 | 88.823 | 1 | 4 | 2 | 0 |
| 53.740 | 12 | 0 | 0 | 4 | 70.766 | 5 | 3 | 0 | 3 | 89.064 | 4 | 3 | 3 | 2 |
| 54.361 | 1 | 2 | 2 | 1 | 71.780 | 21 | 1 | 0 | 5 | | | | | |
| 54.950 | 27 | 2 | 0 | 3 | 73.713 | 29 | 3 | 1 | 3 | | | | | |

**Figure S8**. X-ray diffraction card No. (82-1406) corresponding to silicon oxide "crist" ($SiO_2$) phase formed due to phase separation induced in G2, and G3 samples.

**Table S3**. shows the crystal size of the mica phase in the G0, G1, G2, and G3 samples after sintering at 900 °C for 2 h

| Sample | B (rad) | Cos θ | | θ | k | Mica crystal size* (nm) |
|---|---|---|---|---|---|---|
| **G0** | 0.01252 | 0.96923 | | 14.2497 | 0.9 | 114.2 |
| **G1** | 0.01765 | 0.96926 | | 14.2427 | 0.9 | 81.0 |
| **G2** | 0.02295 | 0.96914 | | 14.2690 | 0.9 | 62.3 |
| **G3** | 0.02848 | 0.96922 | | 14.2511 | 0.9 | 50.2 |

*Depending on the main peak of d-spacing 3.12 Å

**Table S4**. Shows the degree of crystallinity of mica phase in the G0, G1, G2 and G3 samples after sintering at 900 °C for 2 h

| Sample | I | V | Mica Crystalinity* |
|---|---|---|---|
| **G0** | 44.028 | 3.886 | 91.2% |
| **G1** | 43.267 | 5.9261 | 86.3% |
| **G2** | 57.263 | 8.1923 | 85.7% |
| **G3** | 45.267 | 12.456 | 72.5% |

***Depending on the intensity of the main peak of d-spacing 3.12 Å and the intensity of the hollow between it and the peak of d-spacing 3.21Å

**Table S5**. Shows the fluoroapatite crystal size of G0, G1, G2, and G3 after sintering at 900 °C for 2 h

| Sample | B (rad) | Cos θ | Θ | k | fluoroapatite crystal size (nm) * |
|---|---|---|---|---|---|
| **G0** | **-** | **-** | **-** | **-** | **-** |
| **G3** | **0.0191** | **0.9615** | **15.95** | **0.9** | **75.3** |

*Depending on the main peak of d-spacing 2.80 Å

**Table S6**. Shows the average crystal size of G0, G1, G2, and G3 after sintering at 900 °C for 2 h

| Sample | Average crystal size (nm) |
|---|---|
| **G0** | 223.79 |
| **G1** | 188.05 |
| **G2** | 55.98 |
| **G3** | 49.81 |

**Table S7.** Densities of coated and uncoated glass ceramic samples

| Sample | Density without wax ($g/cm^3$) | Density with wax ($g/cm^3$) |
|---|---|---|
| **G0** | 2.80 ± 0.019 | 2.78 ± 0.018 |
| **G1** | 2.77 ± 0.0085 | 2.75 ± 0.0121 |
| **G2** | 2.74 ± 0.013 | 2.73 ± 0.017 |
| **G3** | 2.70 ± 0.011 | 2.69 ± 0.009 |

**FTIR analysis of glass and glass-ceramic samples after SBF immersion**

FTIR analysis was conducted to investigate the formation of the HCA layer on the surface of glass and glass-ceramic samples following immersion in SBF. The fundamental structure of silicate-based glasses consists of a three-dimensional (3D) network of $SiO_4$ tetrahedra, interconnected at their corners to form a continuous framework.[1] The incorporation of alkali ($R_2O$) and alkaline earth (RO) oxides as network modifiers disrupts this connectivity by converting bridging oxygen (BO) into non-bridging oxygen (NBO). This phenomenon, described by Zachariasen[2] and further supported by Warren and Biscoe,[3] leads to structural changes where mobile ions occupy interstitial spaces, thereby altering the glass network and lowering the glass transition temperature.[4-6]

The FTIR spectra reveal significant structural modifications with increasing alkali oxide ($R_2O$/RO) content. Up to 33.3 mol% $R_2O$, $Q^4$ units ($SiO_4$ tetrahedra with four BOs) progressively transform into $Q^3$ units containing three BOs and one NBO. At 33.3 mol% $R_2O$, the network is predominantly composed of $Q^3$ units, which further transition into $Q^2$ units with increasing alkali content, reaching full conversion at 50 mol% $R_2O$.[7, 8] The absorption bands near 1100 $cm^{-1}$ correspond to the stretching vibrations of a single NBO on a $SiO_4$ tetrahedron, indicative of network modification.[9] The bands around 950 $cm^{-1}$ and 900 $cm^{-1}$ are assigned to Si–$O^-$ stretching in tetrahedral units with two and three NBOs, respectively, while the 850 $cm^{-1}$ band corresponds to monomeric $SiO_4^{4-}$ units.[9] The effect of cation interactions is evident, with $Ca^{2+}$ shifting absorption to ~940 $cm^{-1}$, while $Na^+$ interactions result in a shift to ~1040 $cm^{-1}$.[10]

Upon immersion in SBF, the following sequential reactions occur at the sample surface:[11, 12]

1. **Ion Exchange**: A rapid exchange of $Na^+$ and $Ca^{2+}$ in the glass with $H^+$ from the solution:
   $$Si–O–Na^+ + H^+ + OH^- \rightarrow Si–OH + Na^+ \text{ (solution)} + OH^-$$
2. **Silica Dissolution**: The cleavage of Si–O–Si bonds results in the release of soluble silica as Si–(OH) groups and the formation of surface silanol groups:
   $$Si–O–Si + H_2O \rightarrow Si–OH + OH–Si$$
3. **Silica Layer Formation**: Condensation and re-polymerization of surface silanol groups create a silica-rich layer on the glass surface.
4. **Calcium-Phosphate Deposition**: $Ca^{2+}$ and $PO_4^{3-}$ ions migrate through the silica layer, leading to the formation of a calcium-phosphate (Ca-P) rich layer at the surface.
5. **HCA Crystallization**: The incorporation of $OH^-$ and $CO_3^{2-}$ from the solution facilitates the crystallization of the Ca-P layer into hydroxycarbonate apatite (HCA).

Each of these reaction stages is distinctly observed through characteristic changes in the vibrational modes of the surface species, confirming the bioactivity of the glass and its ability to promote HCA layer formation.

The FTIR spectra of glass samples before and after immersion in SBF (Figures S9 and S10) indicate distinct structural changes.

Before SBF immersion, G0 and G1 (2.5 wt% $P_2O_5$) show no prominent bands appearing in the 1100–1200 $cm^{-1}$ range, confirming their amorphous nature (Figure 9). With increasing $P_2O_5$ content (5.0 wt% in G2), a band emerges at 1120 $cm^{-1}$, while in G3 (7.5 wt% $P_2O_5$), additional bands at 1120 $cm^{-1}$ and 1190 $cm^{-1}$ appear, signifying the formation of silicate phases. A broad hump around 1000 $cm^{-1}$ in G0 and G2 sharpens into a distinct band at 1030 $cm^{-1}$ in G2 and

G3, further indicating structural modifications. The functional groups in unreacted glass samples are summarized in Table S9, and the vibration modes of functional groups are included in Table S10.

**Table S9**. Functional groups of the unreacted glass samples before immersion in SBF

| **Peak Range ($cm^{-1}$)** | **G0** | **G1** | **G2** | **G3** |
|---|---|---|---|---|
| 415-540 | Si-O-Si bending vibration (Si-O-Si rocking vibration) | | | |
| | 450, 520 | 415, 480 | 450, 530 | 450, 530 |
| 700-1175 | Si-O-Si anti-symmetric stretching within the $SiO_4$ tetrahedral | | | |
| | 730, 1000 | 750, 1026 | 760, 1026, 1120 | 760, 1026,1120, 1190 |
| 860-940 | Si-O-2NBO | | | |
| | 930 | 940 | 940 | 940 |
| 1400-1550 | C-O(S) | | | |
| | -- | -- | 1450 | 1450 |
| 1600-1650 | H-O-H | | | |
| | 1640 | 1600 | 1610 | 1610 |

**Table S10**. Infrared wavenumber of functional groups on glass samples after SBF immersion

| **Wavenumber ($cm^{-1}$)[9, 13-17]** | **Band** | **Vibrational mode** |
|---|---|---|
| 1080-1350 | P=O | Stretch |
| 800-890 | C-O | Stretch out |
| 700-1175 | Si-O-Si | Tetrahedral |
| 560-610 | P-O | Bend/crystal |
| 550-560 | P-O | Bend/glass |
| 510-530 | P-O | Bend/crystal |
| 450-500 | P-O | Bend/crystal |
| 415-540 | Si-O-Si | Bend |
| 1400-1530 | C-O | stretch |
| 860-1040 | Si-O-(n)NBO | stretch |

After SBF exposure, distinct spectral changes occur due to alkali-hydrogen ion exchange, network dissolution, and the formation of new chemical phases (Figure S10). These transformations are marked by a reduction in the intensity of Si-O-NBO (non-bridging oxygen) modes, which are gradually replaced by Si-OH groups. Over time, the Si-2NBO vibrations are substituted by Si-O-Si bonds, signifying the development of a silica-rich layer. This transition is confirmed by the appearance of Si-O-Si[T] vibrations at specific wavenumbers: 1230 $cm^{-1}$ (G0), 1265 $cm^{-1}$ (G1), 1245 $cm^{-1}$ (G2), and 1235 $cm^{-1}$ (G3). The formation of this silica gel layer in Stage 3 provides nucleation sites for a calcium-phosphate-rich phase, which subsequently crystallizes into hydroxycarbonate apatite (HCA), marking Stages 4 and 5 of bioactivity.

In the G0 sample, the emergence of new bands at 470 $cm^{-1}$, ~560 $cm^{-1}$, and ~1060 $cm^{-1}$ corresponds to P-O bending vibrations, while the appearance of a single band at 890 $cm^{-1}$ is attributed to C-O out-of-plane bending of the carbonate group. A shift in the Si-O-Si(T) band from ~730 $cm^{-1}$ to ~705 $cm^{-1}$, along with the decreased intensity or disappearance of silica bands at 415 $cm^{-1}$, 480 $cm^{-1}$, and 1000 $cm^{-1}$, indicates silicon ion release into the solution. Similarly, in the G1 sample, new bands at ~470 $cm^{-1}$, ~560 $cm^{-1}$, and ~1060 $cm^{-1}$ confirm phosphate formation, while carbonate-related bands appear at 890 $cm^{-1}$ and 1430 $cm^{-1}$. The Si-O-Si(T) vibration shifts from ~750 $cm^{-1}$ to ~725 $cm^{-1}$, with notable reductions in the silica bands at 415 $cm^{-1}$, 480 $cm^{-1}$, and 1026 $cm^{-1}$.

For the G2 sample, new bands at 460 $cm^{-1}$, ~575 $cm^{-1}$, and ~1060 $cm^{-1}$ further indicate P-O bending vibrations, with carbonate contributions seen at 850 $cm^{-1}$ and the splitting of the 1450 $cm^{-1}$ band into two distinct peaks at 1430 $cm^{-1}$ and 1530 $cm^{-1}$. A complete disappearance of Si-O-Si(T) bands at ~760 $cm^{-1}$ and 1120 $cm^{-1}$, along with reductions in silica bands at 450 $cm^{-1}$, 530 $cm^{-1}$, and 1026 $cm^{-1}$, further confirms silicon leaching into the solution. Finally, in the G3 sample, similar trends are observed, with new P-O bending vibration bands at 490 $cm^{-1}$, ~590 $cm^{-1}$, and ~1060 $cm^{-1}$, as well as carbonate-related bands at 890 $cm^{-1}$ and the split 1450 $cm^{-1}$ peak into 1430 $cm^{-1}$ and 1530 $cm^{-1}$. The complete disappearance of Si-O-Si(T) vibrations at ~760 $cm^{-1}$, 1120 $cm^{-1}$, and 1190 $cm^{-1}$, along with reductions in silica bands at 450 $cm^{-1}$, 530 $cm^{-1}$, and 1026 $cm^{-1}$, indicates extensive chemical modification due to prolonged SBF exposure.

Overall, these FTIR spectral changes confirm the progressive transformation of the glass surfaces, demonstrating the sequential stages of silica network dissolution, bioactive silica gel layer formation, and subsequent calcium-phosphate precipitation, ultimately leading to the development of an HCA layer. This suggests that $P_2O_5$ plays a crucial role in the nucleation and growth of HCA, accelerating apatite formation and confirming the bioactivity of $P_2O_5$-containing glasses. In contrast, apatite formation in $P_2O_5$-free glass compositions is significantly slower, reinforcing the importance of $P_2O_5$ in promoting bioactive behavior. These findings highlight the structural dependence of bioactive glass on $P_2O_5$ concentration, providing insight into optimizing compositions for biomedical applications.

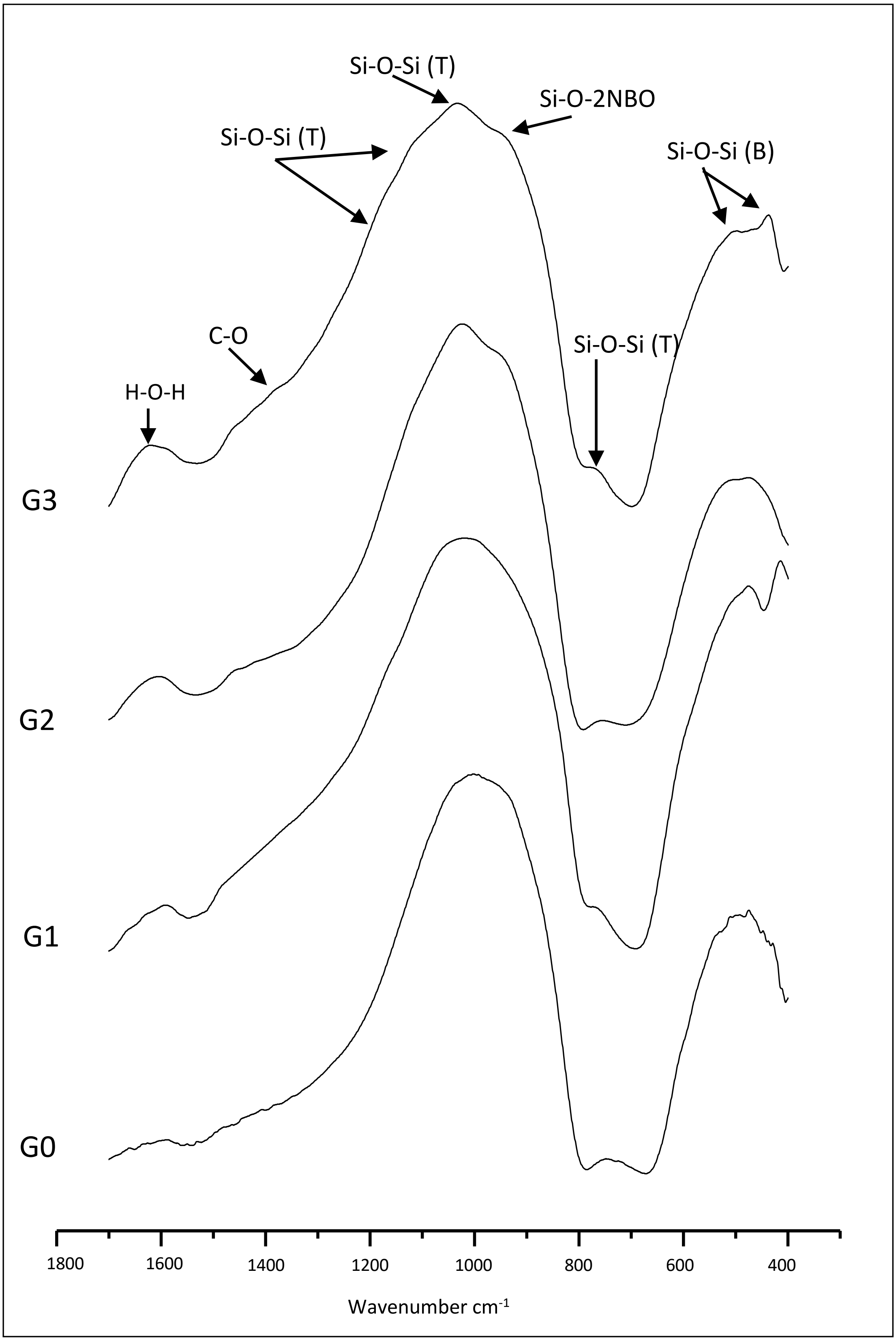


**Figure S9**. FTIR absorbance spectra of glass samples (G0, G1, G2, and G3) before SBF immersion.

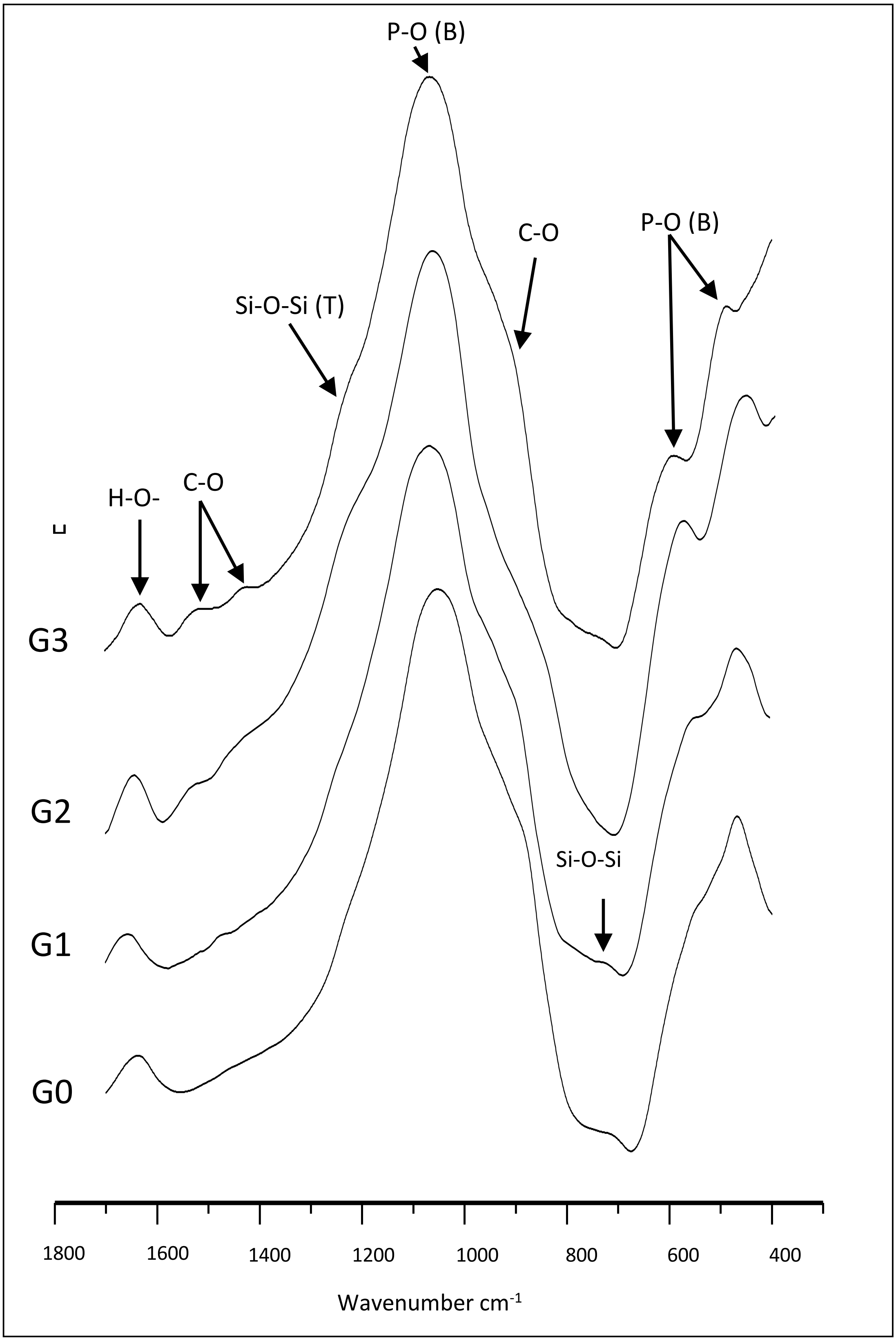


**Figure S10**. FTIR absorbance spectra of glass samples (G0, G1, G2, and G3) after soaking in SBF for 30 days

The FTIR spectra of the glass-ceramic samples heat-treated at 900°C are presented in Figure S11. In the wavelength range of 400-700 $cm^{-1}$, the G0 sample exhibits a sharp band at 418 $cm^{-1}$ and 630 $cm^{-1}$, along with a shoulder at 675 $cm^{-1}$, which can be attributed to Si-O-Si bending vibrations. The incorporation of $P_2O_5$ (2.5 and 5 wt.%) into the base glass (G0) alters these spectral features, transforming the band at 675 $cm^{-1}$ into a broad band at 630 $cm^{-1}$ and shifting the shoulder at 418 $cm^{-1}$ to 518 $cm^{-1}$. This shift suggests a decrease in silicate-containing phases and a concurrent increase in phosphate-containing phases.

The emergence of the phosphate phase is confirmed by a new band appearing at 475 $cm^{-1}$ in G1 and G2, attributed to P-O bending vibrations. Additionally, a broad band at 612 $cm^{-1}$ appears in G2, further indicating phosphate presence. With increasing $P_2O_5$ content in G3 (up to 7.5 wt.%), the Si-O-Si band at 518 $cm^{-1}$ disappears, while the P-O band at 475 $cm^{-1}$ in G1 and G2 splits into two bands at 475 $cm^{-1}$ and 512 $cm^{-1}$ in G3. Similarly, the band at 612 $cm^{-1}$ in G2 splits into a sharp band at 610 $cm^{-1}$ and a shoulder at 575 $cm^{-1}$ in G3, suggesting increased crystallinity of the fluoroapatite phase.

Compared to G0, new bands appear at 1275 $cm^{-1}$ in G1, G2, and G3, attributed to P=O stretching vibrations, further indicating optimal fluoroapatite crystallinity with increasing $P_2O_5$ content. Additionally, in G3, a new band at 1100 $cm^{-1}$, assigned to Si-O-Si stretching vibrations, may be associated with increased crystallinity of enstatite and forsterite phases. These FTIR results align well with the XRD analysis of the glass-ceramic samples heat-treated at 900°C for 2 hours (Figure 4-3b). The functional groups of the unreacted glass-ceramic samples before SBF immersion are summarized in Table S11.

**Table S11**. Functional groups of the unreacted glass-ceramic samples before immersion in SBF

| **Peak Range $cm^{-1}$** | **G0** | **G1** | **G2** | **G3** |
|---|---|---|---|---|
| **415-540** | Si-O-Si bending vibration (Si-O-Si rocking vibration) | | | |
| | 418, 518 | 418, 518 | 418, 518 | 418 |
| **630-680 700-1175** | Si-O-Si anti-symmetric stretching within $SiO_4$ tetrahedral (123) | | | |
| | 630, 680, 1175 | 630, 680, 1066 | 630, 680, 1057 | 674, 705, 1050 |
| **1080-1380** | P = O stretching vibration. | | | |
| | -- | 1275 | 1280 | 1100, 1276 |
| **860-990** | Si-O-(n)NBO | | | |
| | 880, 975 | 882, 980 | 882, 980 | 882, 990 |
| **510-530 550-615** | P-O bending vibration | | | |
| | -- | 475 | 475, 612 | 475,512, 575, 610 |
| **1400-1550** | C-O(S) | | | |
| | -- | 1400, 1455 | 1457 | 1429 |
| **1600-1650** | H-O-H | | | |
| | -- | 1630 | 1640 | 1645 |

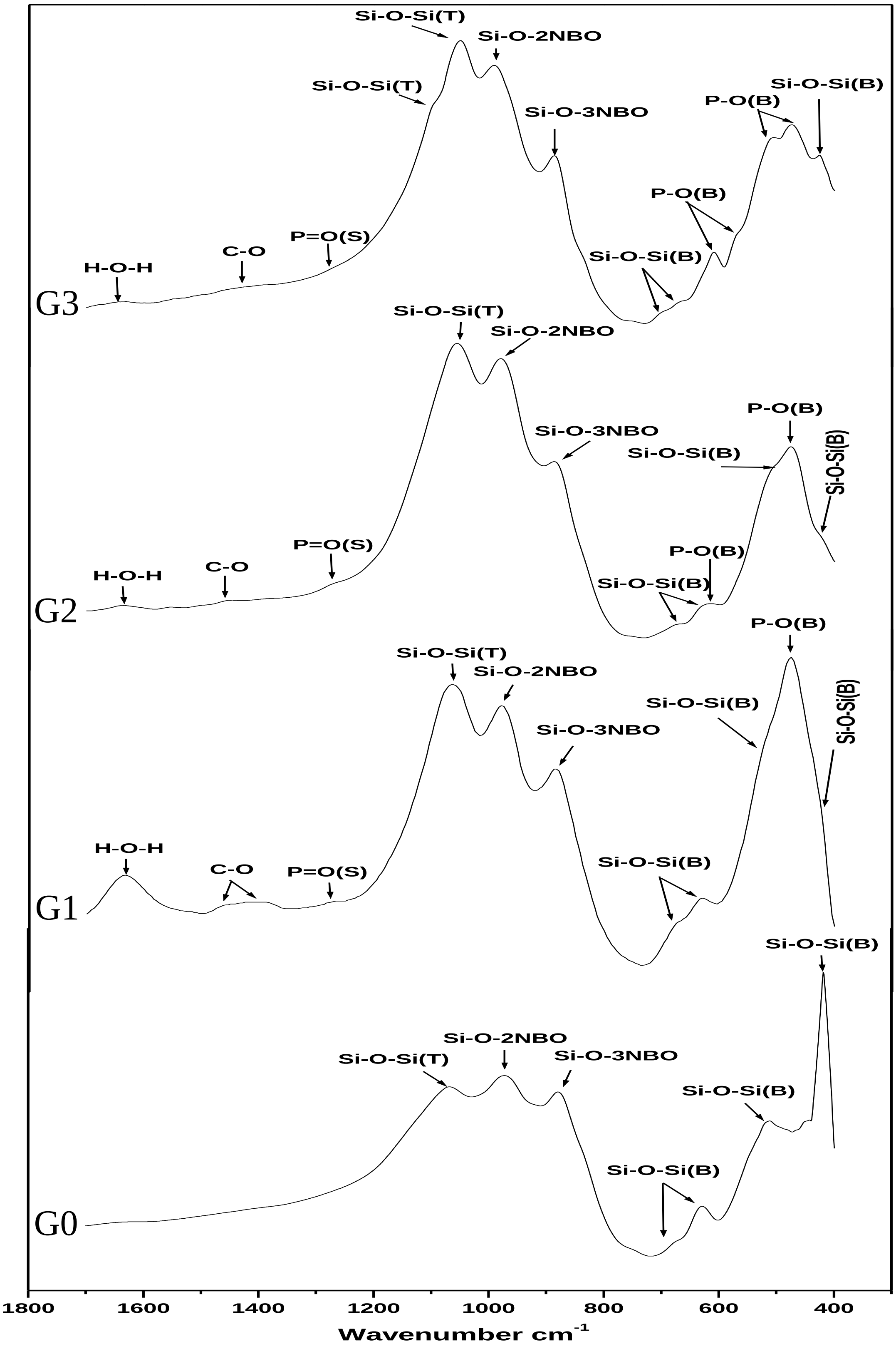


**Figure S11**. FTIR absorbance spectra for glass-ceramic samples (G0, G1, G2, and G3) before SBF immersion.

After immersion in SBF (Figure S12), all reaction stages are represented by changes in the vibration modes of the chemical species of the powder. The alkali-hydrogen ion exchange and network dissolution (Stages 1 and 2) rapidly reduce the intensity of the Si-O-NBO modes and replace them with Si-OH. However, because the sample was examined after 30 days of immersion in SBF, it was difficult to follow these stages. As the reactions continue, the two non-bridging oxygen (Si-2NBO) modes disappear in the G0 sample, while their intensity is reduced in G1, G2, and G3 samples, being replaced by a new mode assigned to the Si-O-Si bond vibration between adjacent $SiO_4$ tetrahedra. Meanwhile, the bands assigned to three non-bridging oxygen (Si-3NBO) modes are replaced with Si-OH and increase in intensity.

The presence of the $SiO_2$-rich layer is recognized by the appearance of the Si-O-Si [T] vibration, which appears as a shoulder at 1240 $cm^{-1}$, 1100 $cm^{-1}$, 1235 $cm^{-1}$, and 1240 $cm^{-1}$. This vibration mode corresponds to the formation of the silica gel layer through Stage 3 (polycondensation reaction between neighboring surface silanol groups), which acts as a nucleation site for a calcium-phosphate-rich layer that can crystallize into hydroxycarbonate apatite (HCA). The formation of the HCA layer is evident from the reduction of silica glass bands and the development of HCA bands (Stages 4 and 5).

In the G0 sample, new bands at ~470 $cm^{-1}$ and ~1046 $cm^{-1}$ are attributed to vibrational modes of P-O bending, and a weak band at ~1315 $cm^{-1}$ corresponds to P=O stretching vibrations. Additionally, new bands at ~1390 $cm^{-1}$ and ~1465 $cm^{-1}$ are assigned to C-O stretching vibrations. The Si-O-Si (B) bands at ~418 $cm^{-1}$, ~518 $cm^{-1}$, ~630 $cm^{-1}$, and ~680 $cm^{-1}$ show reduced intensity due to the release of silicon ions into the solution. The Si-O-2NBO band at ~975 cm-1 disappears completely, while the Si-O-3NBO band is replaced by Si-OH due to weak ion exchange, as confirmed by UV analysis (Figure 8).

In the G1 sample, the P-O (B) band at 475 $cm^{-1}$ splits into two bands at 475 $cm^{-1}$ and 512 $cm^{-1}$, while a new band at 1035 $cm^{-1}$ indicates increased apatite formation. The band at 1400 $cm^{-1}$ splits into two weaker bands at 1388 $cm^{-1}$ and 1425 $cm^{-1}$, assigned to C-O stretching vibration. The Si-O-Si (B) band at ~518 $cm^{-1}$ disappears completely, and its intensity at 630 $cm^{-1}$ and 680 $cm^{-1}$ decreases due to silicon ion release. The Si-O-2NBO peak at 980 cm-1 shifts to a shoulder at ~960 $cm^{-1}$, while the Si-O-3NBO band at 882 $cm^{-1}$ is replaced by Si-OH due to weak ion exchange.

In the G2 sample, the P-O (B) band at 475 $cm^{-1}$ splits into a clear band at 475 $cm^{-1}$ and a weak band at 512 $cm^{-1}$. The band at 1457 $cm^{-1}$ splits into two weaker bands at 1450 $cm^{-1}$ and 1550 $cm^{-1}$, assigned to C-O stretching vibration. The Si-O-Si (B) band at ~518 $cm^{-1}$ disappears completely, and its intensity at 630 $cm^{-1}$ and 680 $cm^{-1}$ decreases due to silicon ion release. The Si-O-2NBO peak at 980 cm-1 becomes a shoulder at ~980 $cm^{-1}$, while the Si-O-3NBO band at 882 $cm^{-1}$ is replaced by Si-OH due to weak ion exchange.

In the G3 sample, there is only a slight increase in the intensity of the 512 $cm^{-1}$ band assigned to P-O (B), indicating a minimal amount of apatite formation on the surface. Similarly, the intensity of the 1445 $cm^{-1}$ band shows only a slight increase, suggesting limited carbonate formation.

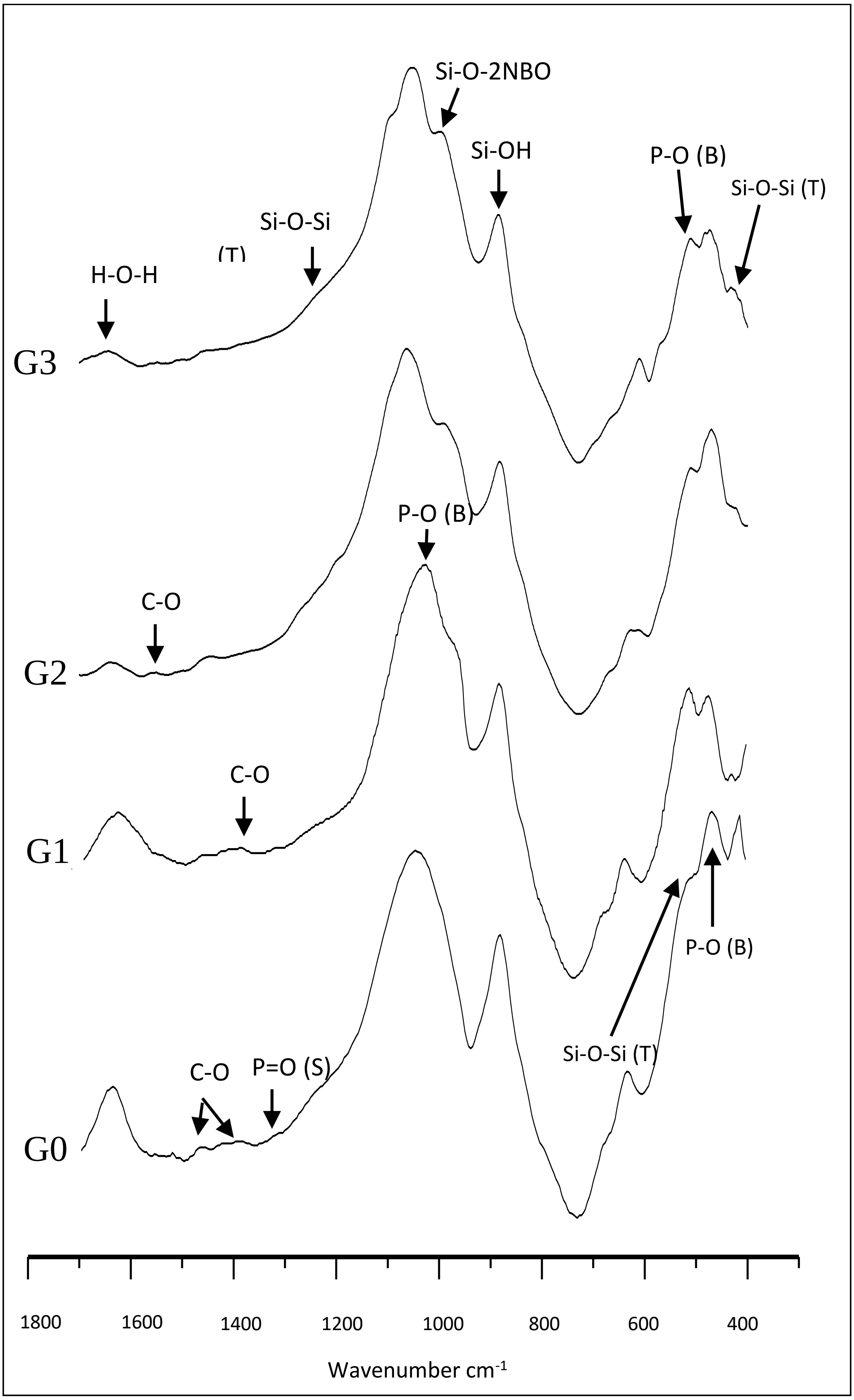


**Figure S12.** FTIR absorbance spectra for glass-ceramic samples (G0, G1, G2, and G3) after soaking in SBF for 30 days
